# Investigation of Human Endogenous Retrovirus-K (ERVK) Expression and Function in Normal Placentation and Preterm Pregnancy

**DOI:** 10.1101/2021.11.22.469425

**Authors:** Jimi L. Rosenkrantz, Michael Martinez, Adithi Mahankali, Lucia Carbone, Shawn L. Chavez

## Abstract

**Background:** There is a growing body of evidence indicating the importance of endogenous retrovirus (ERV) derived proteins during early development and reproduction in mammals. Recently, a protein derived from the youngest ERV in humans, ERVK (HML2), was shown to be expressed during human placentation. Since a number of highly similar ERVK proviral loci exist across the human genome, locus-specific analysis of ERVK transcription and identification of the coding sequence expressed in the human placenta is difficult. Thus, despite its activity in early human development, the native expression and function of ERVK in the human placenta remains largely uncharacterized.

**Results:** In this study, we comprehensively examined locus-specific ERVK transcription across several human placental tissues and cell types. Through a combination of RNA-seq and siRNA knock-down analyses, we identified the expression of a single ERVK locus, ERVK11q23.3, as (1) being significantly upregulated in preterm compared to term placenta, (2) predominantly expressed by mononuclear trophoblasts, (3) capable of encoding a truncated viral-like envelope protein, and (4) contributing to the expression cytokines involved in both antiviral and anti-inflammatory innate immune responses in human placental trophoblasts and BeWo choriocarcinoma cells, respectively.

**Conclusions:** Collectively, the results of this study highlight the utility of studying locus-specific ERVK expression, provide a thorough characterization of locus-specific ERVK transcription from human placental tissues, and indicate that altered expression of placental ERVK11q23.3 influences interferon antiviral response, which may contribute to preterm birth and other pregnancy complications.

## Background

Mammalian genomes are littered with thousands of copies of endogenous retroviruses (ERVs), mobile genetic elements that are relics of ancient retroviral infections. ERVs are abundant within the DNA of all vertebrates and specifically comprise ∼8% of the human genome [1]. Despite being considered ‘junk DNA’ for many years, there has been a growing interest in the biological role of ERV-derived proteins in early human development and reproduction, including placentation. ERV proviral insertions within the human genome are classified into different groups based on sequence similarity [2]. Full-length ERV insertions possess similar genomic organization to exogenous retroviruses, including two flanking long terminal repeats (LTRs), an internal sequence corresponding to group-specific antigen (*gag)*, protease (*pro*), polymerase (*pol),* and envelope (*env)* proviral genes. The vast majority of ERVs within the genome are truncated and/or have become highly mutated and thus, are unable to produce functional viral proteins and become infectious [3]. However, a few ERV insertions have remained relatively well-conserved, and contain open reading frames (ORFs) that can encode viral-like proteins. Several ERV viral-like proteins expressed during development are known to play important physiological roles. The most notable examples of this are the Syncytin proteins, which are expressed during human placentation. Syncytin-1 [4] and Syncytin-2 [5] are encoded from *env* genes of ERVW and ERVFRD groups, respectively. Similar to exogenous retroviral envelope proteins, Syncytin proteins contain fusion peptide (FP) and/or immunosuppression (ISU) domains. Exogenous retroviruses utilize the FP and ISU domain to facilitate fusion of the viral particle into the host cell and suppression of the host immune response, respectively [6, 7]. Unlike exogenous retroviruses, the Syncytin proteins, which contain one or both of these domains, have been co-opted to facilitate important processes underlying normal human placentation. Specifically, Syncytin FPs are utilized by mononuclear cytotrophoblasts (CTBs) to facilitate the cell fusion underlying the formation and maintenance of the multinucleated syncytiotrophoblast (STB) layer [5,8,9], and there is evidence suggesting that the ISU domain of Syncytin-2 aids maternal immune evasion during pregnancy [10].

Similar to Syncytins, ERVK envelope (ERVK-env) protein has been shown to be expressed during human placentation [11]. This protein is derived from the ERVK (HML-2) group, the youngest and most recently expanded ERV in primate genomes [12–15]. Thus, unlike the ERV families encoding the Syncytin proteins, dozens of ERVK proviral insertions with intact envelope ORFs exist across the human genome. While there are no replication-competent ERVK proviral insertions, the presence of polymorphic ERVK insertions in human indicates that the family was active and infectious up until at least 5-6 million years ago [16, 17]. A consequence of this recent activity is that many ERVK loci are highly-similar, which makes it difficult to analyze locus-specific ERVK expression using traditional short-read RNA-seq data. Due to these challenges, a thorough characterization ERVK placental expression, including locus-specific transcription, splicing, and putative protein-coding sequences has been lacking.

Several studies using ERVK ancestral-predicted consensus sequences have identified a functional FP [18–20] and ISU domain [21] within the ERVK-env protein, and its ability to elicit cell fusion when ectopically expressed in cell lines or incorporated into viral particles [18–20], as well as inhibit immune cell proliferation *in vitro* [21]. These reports suggest that placental ERVK expression, and specifically ERVK-env protein expression, may facilitate trophoblast cell fusion and/or maternal immunosuppression during normal placentation. However, since it is unclear which ERVK loci are expressed in the placenta, whether ERVK-env is fusogenic and/or immunosuppressive at this site remains unknown. We hypothesized that ERVK expression, and specifically ERVK-env protein, has been co-opted to facilitate trophoblast cell fusion and/or maternal immunosuppression during normal placentation. Additionally, since trophoblast dysfunction and heightened inflammation are associated with pregnancy complications [22, 23], we further postulate that abnormal ERVK expression is associated with placental dysfunction in preterm birth. Thus, to test these hypotheses, we aimed to (1) thoroughly characterize placental ERVK expression, including locus-specific transcription levels and ERVK-env protein-coding potential, (2) identify ERVK loci differentially expressed between placenta from healthy and pathological pregnancies, and (3) assess the fusogenic and/or immunomodulatory function of placentally-expressed ERVK.

## Results

### ERVK-env protein is expressed in the STB layer of term placenta and its expression varies across preterm placentas

ERVK-env protein expression has previously been documented in human placental tissues from healthy pregnancies [11]. Since functional FP and ISU domains were identified within the ancestral-predicted ERVK-env protein and both trophoblast dysfunction and heightened inflammation are commonly associated with preeclampsia and preterm birth [22, 23], we hypothesized that aberrant placental ERVK-env protein expression may be associated with these pregnancy complications. Thus, we sought to compare placental ERVK-env protein expression and localization between normal and pathological pregnancy conditions. For this, we preformed immunohistochemical (IHC) staining on human placental tissue collected from term (n=4) and preterm (n=4) pregnancies using a well-established commercially available anti-ERVK-env monoclonal antibody [11] (**Table 1**). In term placenta samples, ERVK-env staining was observed at the maternal-fetal interface in multinucleated STBs. The staining pattern was diffuse throughout the cytoplasm and microvilli of the STB (**Figure 1A**), analogous to the known STB secreted proteins, CGB and KISS1 [24]. A similar expression pattern was observed in preterm placental tissue; however, variable staining intensities were noted across the four different samples examined (**Figure 1A**). The absence of non-specific staining was confirmed by substituting the anti-ERVK-env antibody with an appropriate mouse igG2A isotype control (**Figure 1B**). Collectively, these results show that ERVK-env protein is expressed by placental trophoblasts, and suggest that the expression level may be affected in placenta from preterm birth pregnancies.

**Figure 1.**
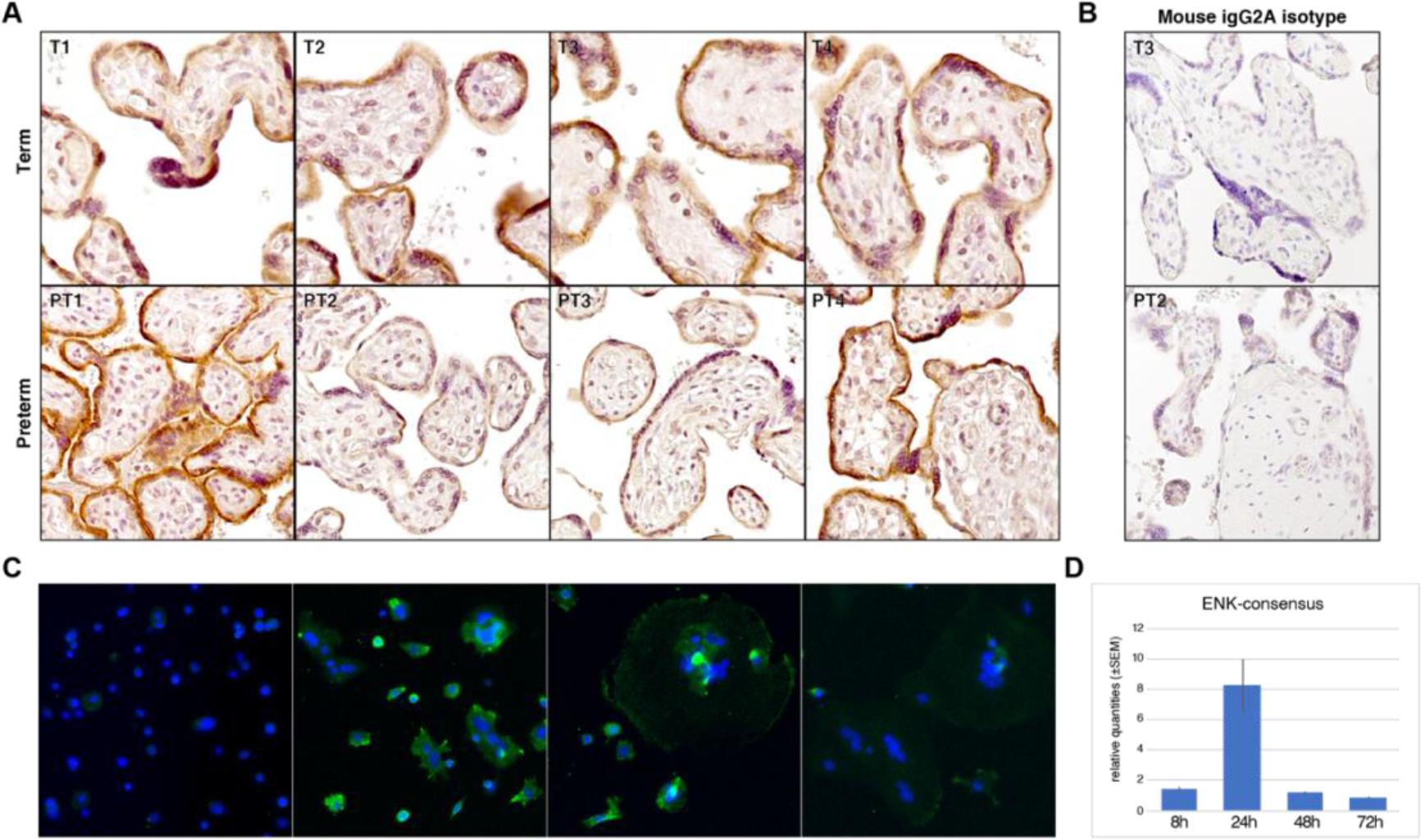
ERVK-env protein expression in placentas and isolated trophoblasts. (A) IHC staining of human term (n=4) and preterm (n=4) placental tissue sections for ERVK-env TM protein and (B) mouse igG2A isotype negative control. Positive staining (DAB, brown), nuclear counterstain (purple). (C) ERVK-env IF staining of HPTs at 8, 24, 48, 72 h (left to right); ERVK-env (green) and DAPI nuclear counterstain (blue). (D) qRT-PCR of HPTs at 8, 24, 48, 72 h (n=3 each); four technical replicates were used per sample and samples were normalized to GAPDH.

**Table 1.** IHC tissue details.

| ID | Category | Fetus sex | Gest Age (wks.) | Labor | C-section | Mat Age (yr.) | Mat Race |
| --- | --- | --- | --- | --- | --- | --- | --- |
| PT1 | Preterm | Female | 35 | No | Yes | 24 | White Hispanic |
| PT2 | Preterm | Female | 35 | No | Yes | 23 | White Non-Hispanic |
| PT3 | Preterm | Female | 35 | No | Yes | 19 | White Non-Hispanic |
| PT4 | Preterm | Male | 35 | No | Yes | 22 | White Hispanic |
| T1 | Term | Female | 39 | No | Yes | 28 | White Non-Hispanic |
| T2 | Term | Male | 39 | No | Yes | 32 | White Non-Hispanic |
| T3 | Term | Male | 39 | No | Yes | 24 | White Non-Hispanic |
| T4 | Term | Male | 39 | No | Yes | 29 | White Non-Hispanic |

### ERVK-env protein is predominantly expressed in unfused mononuclear CTBs *in vitro*

Mononuclear CTB cell fusion is critical for the formation and maintenance of the multinucleated STB layer in the placenta [25, 26]. Similar to the Syncytin proteins, the ERVK-env protein is speculated to play a role in CTB fusion and STB formation. This is largely based on the fusogenic capabilities of ancestral-predicted recombinant ERVK-env proteins [18–20], and the report that native ERVK-env protein is expressed at the cell surface of villous CTBs in the human placenta [11], which is the expected location of a protein involved in CTB cell-cell fusion [27]. To further evaluate native ERVK-env expression and localization throughout CTB differentiation and fusion, we utilized human primary trophoblast (HPT) cell cultures, which consist of highly purified CTBs that spontaneously differentiate and fuse into STBs over time [25, 26]. Thus, we examined ERVK-env protein and mRNA expression in HPTs after 8, 24, 48, and 72 h in culture (**Figure 1C-D**). ERVK-env protein expression and localization was assessed via immunofluorescence (IF) using a well-established monoclonal antibody targeting the TM envelope protein of ERVK (anti-ENK); while, ERVK-env RNA expression was assessed via qRT-PCR using primers designed to amplify the ancestral-predicted ERVK-env sequence (ENK-consensus) [28]. Consistent with previous reports [11], the IF results showed that ERVK-env protein expression appeared most prominent around the membrane of mononuclear CTBs 24 h after culturing (**Figure 1C**). Cytoplasmic staining of multinucleated STBs was also observed in HPT cultures, supporting the findings from our IHC analysis of bulk placental tissues (**Figure 1A,C**). The qRT-PCR results further demonstrated that the highest ERVK-env transcription levels were detected at 24 h of culture (**Figure 1D**). Collectively, these data indicate that ERVK-env expression peaks after 24 h in culture and that its protein is predominantly located at the plasma membrane of unfused mononuclear CTBs, which is consistent with ERVK being involved in trophoblast fusion.

### Multiple ERVK loci with intact envelope ORFs are transcribed in term and preterm placentas

A number of ERVK proviral loci exist across the human genome [29], and the transcriptional activity of each locus is dependent on the tissue/cell type [30]. To identify which specific ERVK proviral loci are expressed in the placenta, we performed locus-specific ERVK transcriptional analysis using RNA-seq data from term (n=5) and preterm (n=5) bulk placental samples (**Table 2**). Since there are a number of highly similar ERVK loci in the human genome, genomic regions around ERVK loci are often shared and exhibit low mappability. Thus, non-uniquely mapped RNA-seq reads were discarded from this analysis to avoid non-specific ERVK locus expression. Approximately 49% (61/124) of all ERVK loci (**Table 3**) were expressed (mean normalized read count >1) in the placental tissues (**Figure 2**) and a number of these were also predicted to contain envelope ORFs possessing the monoclonal antibody epitope [11]. This includes ERVK loci with predicted full-length (>588aa) envelope ORFs (ERVK6q25.1, ERVK1q21.3a, ERVK6p21.1, ERVK1q22, ERVK1p31.1a, ERVK1p34.3) [29]; and those with partial (>300aa) envelope ORFs (ERVK3q21.2b, ERVK1q21.3b, ERVK1q23.3, ERVK4q35.2, ERVK5q33.2, ERVKXq28b) [29]. Several other ERVK loci containing partial envelope ORFs with a putative Furin cleavage site [31], but did not contain the monoclonal antibody epitope [31]. Because the Furin cleavage site is necessary for the formation of a fusogenic TM protein [31], expression from these loci may result in functional Furin-cleaved ERVK-env proteins that are undetectable by the monoclonal antibody. Nevertheless, the immunostaining results in **Figure 1** likely reflect envelope protein expression from one or more of the ERVK loci with full-length or partial envelope ORFs that contained both a Furin cleavage site and the epitope of the antibody used in this study.

**Figure 2.**
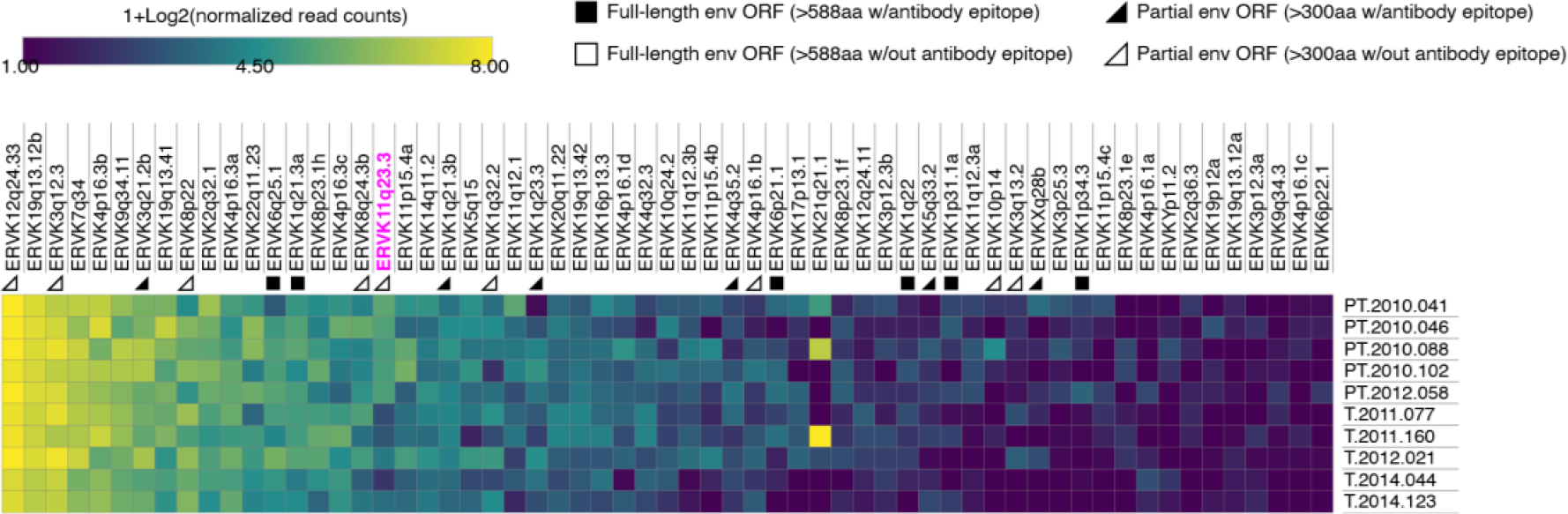
ERVK locus-specific RNA expression analysis in term and preterm placentas. Heatmap depicting ERVK locus-specific RNA expression levels in human term (n=5) and preterm (n=5) placental tissue samples. Only loci with mean normalized read count >1 are shown (n=61). Loci with full-length envelope ORFs (>588aa, square), partial envelope ORF (>300aa, triangle), containing ERVK-env antibody epitope (black), or not containing ERVK-env antibody epitope (white) are shown below each locus.

**Table 2.**
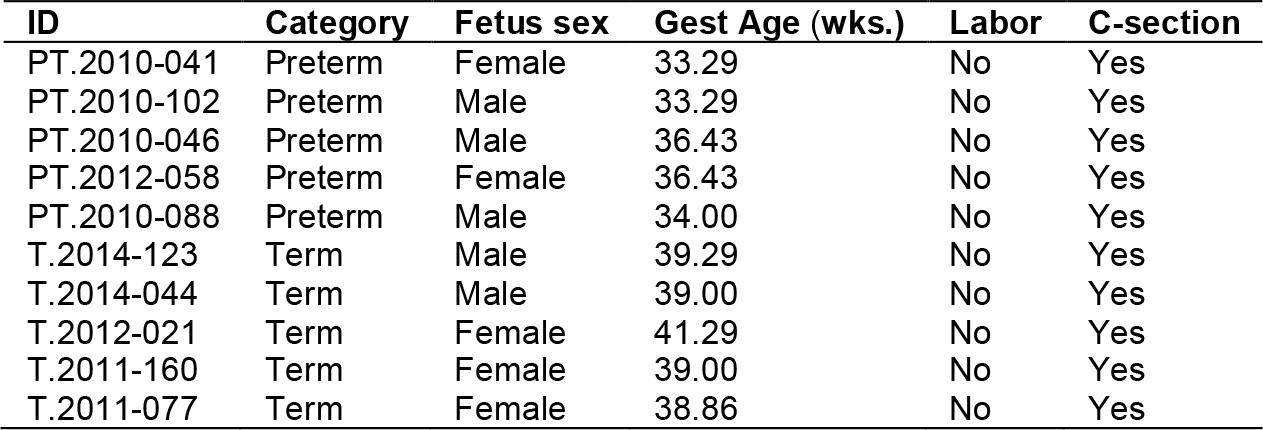
RNA-seq sample details.

**Table 3.** Details of ERVK loci (n=124)

| ID | chr | Start | stop | strand | Longest env ORF (aa) | Antibody epitope | ENK-consensus primer |
| --- | --- | --- | --- | --- | --- | --- | --- |
| ERVK1p36.21a | chr1 | 12780114 | 12785935 | - |  |  |  |
| ERVK1p36.21b | chr1 | 13012446 | 13021988 | - | 131 |  |  |
| ERVK1p36.21c | chr1 | 13206971 | 13216513 | + | 296 |  |  |
| ERVK1p36.21d | chr1 | 13352735 | 13362257 | + | 162 | yes |  |
| ERVK1p35.1 | chr1 | 33063515 | 33065110 | + |  |  |  |
| ERVK1p34.3 | chr1 | 36488983 | 36491127 | - | 588 | yes |  |
| ERVK1p31.1a | chr1 | 75377085 | 75383458 | + | 588 | yes | yes |
| ERVK1p31.1b | chr1 | 82223389 | 82223522 | - |  |  |  |
| ERVK1q21.3a | chr1 | 150632807 | 150635885 | + | 588 | yes |  |
| ERVK1q21.3b | chr1 | 150632885 | 150634662 | - | 542 | yes |  |
| ERVK1q22 | chr1 | 155626665 | 155635845 | - | 1375 | yes | yes |
| ERVK1q23.3 | chr1 | 160691753 | 160698953 | + | 542 | yes | yes |
| ERVK1q24.1 | chr1 | 166605365 | 166611021 | - | 309 |  |  |
| ERVK1q32.2 | chr1 | 207635111 | 207639291 | - | 309 |  |  |
| ERVK1q43 | chr1 | 238762294 | 238764473 | - | 588 | yes |  |
| ERVK2q21.1 | chr2 | 129961964 | 129965044 | - | 687 | yes | yes |
| ERVK2q32.1 | chr2 | 186520906 | 186522372 | + | 170 |  |  |
| ERVK2q36.3 | chr2 | 229180638 | 229180771 | - |  |  |  |
| ERVK3p26.1 | chr3 | 7061961 | 7062102 | + |  |  |  |
| ERVK3p25.3 | chr3 | 9847661 | 9854552 | - | 113 |  |  |
| ERVK3p12.3a | chr3 | 75536829 | 75538747 | + |  |  |  |
| ERVK3p12.3b | chr3 | 75551313 | 75559999 | + | 245 |  |  |
| ERVK3q12.3 | chr3 | 101691892 | 101701015 | + | 416 |  | yes |
| ERVK3q13.2 | chr3 | 113024276 | 113033435 | - | 597 (pol/env) |  | yes |
| ERVK3q21.2a | chr3 | 125799250 | 125801161 | - |  |  |  |
| ERVK3q21.2b | chr3 | 125890458 | 125899596 | + | 312 | yes | yes |
| ERVK3q22.1 | chr3 | 130211771 | 130212782 | + |  |  |  |
| ERVK3q24 | chr3 | 148563689 | 148567609 | - |  |  |  |
| ERVK3q27.2 | chr3 | 185562547 | 185571727 | - | 312 | yes |  |
| ERVK4p16.3a | chr4 | 241199 | 245565 | + |  |  |  |
| ERVK4p16.3b | chr4 | 3977323 | 3986912 | - | 234 |  |  |
| ERVK4p16.3c | chr4 | 4073396 | 4075313 | + |  |  |  |
| ERVK4p16.1a | chr4 | 9034416 | 9036346 | - |  |  |  |
| ERVK4p16.1b | chr4 | 9121785 | 9131367 | + | 342 |  |  |
| ERVK4p16.1c | chr4 | 9567313 | 9569224 | - |  |  |  |
| ERVK4p16.1d | chr4 | 9657955 | 9667550 | + | 234 |  |  |
| ERVK4q13.2 | chr4 | 68597990 | 68603505 | + | 113 | yes |  |
| ERVK4q32.1 | chr4 | 160658785 | 160661208 | + |  |  |  |
| ERVK4q32.3 | chr4 | 164995687 | 165002916 | + | 192 |  |  |
| ERVK4q35.2 | chr4 | 190106258 | 190113546 | - | 312 | yes |  |
| ERVK5p13.3 | chr5 | 30486652 | 30496098 | - | 312 | yes |  |
| ERVK5p12 | chr5 | 46000056 | 46009900 | - | 140 |  |  |
| ERVK5q15 | chr5 | 93456673 | 93458206 | - |  |  |  |
| ERVK5q31.1 | chr5 | 136413959 | 136414650 | - |  |  |  |
| ERVK5q33.2 | chr5 | 154635952 | 154644655 | - | 312 | yes |  |
| ERVK5q33.3 | chr5 | 156657705 | 156666885 | - | 312 | yes | yes |
| ERVK6p25.2 | chr6 | 3054799 | 3055508 | + |  |  |  |
| ERVK6p22.1 | chr6 | 28682590 | 28692958 | + | 134 |  |  |
| ERVK6p21.1 | chr6 | 42893670 | 42903629 | - | 698 | yes |  |
| ERVK6q11.1 | chr6 | 60654986 | 60660975 | + |  |  |  |
| ERVK6q13 | chr6 | 73333257 | 73333407 | - | 162 | yes |  |
| ERVK6q14.1 | chr6 | 77716944 | 77726366 | - | 698 | yes | yes |
| ERVK6q25.1 | chr6 | 150859612 | 150862438 | + | 699 | yes |  |
| ERVK7p22.1 | chr7 | 4582425 | 4600400 | - | 699 | yes | yes |
| ERVK7p14.3 | chr7 | 30718255 | 30719589 | - |  |  |  |
| ERVK7q11.21 | chr7 | 66004683 | 66007803 | - | 150 |  |  |
| ERVK7q22.2 | chr7 | 104748901 | 104752819 | - |  |  |  |
| ERVK7q34 | chr7 | 141752118 | 141756120 | - |  |  |  |
| ERVK8p23.1a | chr8 | 7126986 | 7128897 | - |  |  |  |
| ERVK8p23.1b | chr8 | 7185490 | 7187404 | + |  |  |  |
| ERVK8p23.1c | chr8 | 7497874 | 7507337 | - | 699 | yes | yes |
| ERVK8p23.1d | chr8 | 8100097 | 8102011 | - |  |  |  |
| ERVK8p23.1e | chr8 | 8197177 | 8206699 | + | 264 |  |  |
| ERVK8p23.1f | chr8 | 12216460 | 12225988 | - | 126 |  |  |
| ERVK8p23.1g | chr8 | 12458982 | 12468498 | - | 261 | yes |  |
| ERVK8p23.1h | chr8 | 12565086 | 12567003 | + |  |  |  |
| ERVK8p23.1i | chr8 | 12623205 | 12625104 | + |  |  |  |
| ERVK8p22 | chr8 | 17908095 | 17916431 | - | 379 |  |  |
| ERVK8q11.1 | chr8 | 46264027 | 46272039 | - | 264 |  |  |
| ERVK8q24.3a | chr8 | 139459905 | 139462993 | - | 699 | yes | yes |
| ERVK8q24.3b | chr8 | 145021243 | 145028834 | - | 309 |  |  |
| ERVK9q34.11 | chr9 | 128850235 | 128857457 | + | 60 |  |  |
| ERVK9q34.3 | chr9 | 136780313 | 136789776 | - | 79 | yes |  |
| ERVK10p14 | chr10 | 6824178 | 6833641 | - | 322 |  |  |
| ERVK10q22.3 | chr10 | 80108250 | 80108670 | + |  |  |  |
| ERVK10q24.2 | chr10 | 99820811 | 99827959 | - | 214 |  |  |
| ERVK11p15.4a | chr11 | 3447425 | 3456979 | - | 164 |  |  |
| ERVK11p15.4b | chr11 | 3544465 | 3546375 | + |  |  |  |
| ERVK11p15.4c | chr11 | 4459434 | 4461473 | - |  |  |  |
| ERVK11q12.1 | chr11 | 59000008 | 59005723 | + | 189 |  |  |
| ERVK11q12.3a | chr11 | 62368490 | 62370999 | - | 230 |  |  |
| ERVK11q12.3b | chr11 | 62375544 | 62383091 | - |  |  |  |
| ERVK11q13.2 | chr11 | 67920365 | 67922280 | + |  |  |  |
| ERVK11q13.4a | chr11 | 71673667 | 71675588 | - |  |  |  |
| ERVK11q13.4b | chr11 | 71764073 | 71764819 | + | 162 |  |  |
| ERVK11q22.1 | chr11 | 101695062 | 101704528 | + | 661 |  |  |
| ERVK11q23.3 | chr11 | 118721014 | 118730174 | - | 438 |  | yes |
| ERVK12p11.1 | chr12 | 34619619 | 34629282 | - | 215 |  |  |
| ERVK12q14.1 | chr12 | 58327458 | 58336915 | - | 698 | yes | yes |
| ERVK12q23.3 | chr12 | 107826195 | 107827421 | - |  |  |  |
| ERVK12q24.11 | chr12 | 110570037 | 110571520 | + |  |  |  |
| ERVK12q24.33 | chr12 | 133090535 | 133096478 | - | 358 |  |  |
| ERVK13q12.13 | chr13 | 26509516 | 26510128 | + |  |  |  |
| ERVK14q11.2 | chr14 | 24009695 | 24015776 | - |  |  |  |
| ERVK14q32.33 | chr14 | 105673312 | 105676203 | + |  |  |  |
| ERVK15q25.2 | chr15 | 84160267 | 84163612 | + |  |  |  |
| ERVK16p13.3 | chr16 | 2926158 | 2927660 | + |  |  |  |
| ERVK16p11.2a | chr16 | 34412056 | 34414804 | - | 550 | yes |  |
| ERVK16p11.2b | chr16 | 34997025 | 34999771 | + | 550 | yes |  |
| ERVK16q23.3 | chr16 | 83702132 | 83702178 | - |  |  |  |
| ERVK17p13.1 | chr17 | 8056336 | 8063901 | + |  |  |  |
| ERVK18q21.33 | chr18 | 63851825 | 63852338 | - |  |  |  |
| ERVK19p13.3 | chr19 | 385094 | 387637 | + | 114 |  |  |
| ERVK19p12a | chr19 | 20276590 | 20286703 | + | 285 | yes |  |
| ERVK19p12b | chr19 | 22575021 | 22581759 | + | 312 |  |  |
| ERVK19q11 | chr19 | 27637589 | 27646453 | - | 699 | yes | yes |
| ERVK19q13.12a | chr19 | 35572526 | 35576532 | - | 268 |  | yes |
| ERVK19q13.12b | chr19 | 37106646 | 37116164 | - | 206 |  |  |
| ERVK19q13.41 | chr19 | 52745022 | 52750454 | - | 159 |  |  |
| ERVK19q13.42 | chr19 | 53359094 | 53364791 | + | 215 |  |  |
| ERVK20p11.21a | chr20 | 23693938 | 23695327 | - |  |  |  |
| ERVK20p11.21b | chr20 | 23755866 | 23757247 | - |  |  |  |
| ERVK20q11.22 | chr20 | 34126943 | 34136578 | + | 139 |  |  |
| ERVK21q21.1 | chr21 | 18561340 | 18569644 | - | 55 | yes |  |
| ERVK22q11.21 | chr22 | 18938673 | 18947848 | + | 1171 | yes | yes |
| ERVK22q11.23 | chr22 | 23536061 | 23548428 | + | 141 |  |  |
| ERVK22q13.2 | chr22 | 40696388 | 40696742 | - |  |  |  |
| ERVKXq11.1 | chrX | 62740078 | 62742584 | + |  |  |  |
| ERVKXq12 | chrX | 66464289 | 66466342 | - |  |  |  |
| ERVKXq28a | chrX | 154588661 | 154591282 | + |  |  |  |
| ERVKXq28b | chrX | 154608422 | 154615762 | - | 336 | yes |  |
| ERVKYp11.2 | chrY | 6958399 | 6965343 | - | 225 |  |  |
| ERVKYq11.23a | chrY | 24251689 | 24254888 | - |  |  |  |
| ERVKYq11.23b | chrY | 25415254 | 25418454 | + |  |  |  |

### ERVK11q23.3 RNA expression is upregulated in preterm compared to term placental tissue

To determine whether ERVK loci or other genes were differentially expressed between human term and preterm placentas, we used the RNA-seq data generated from both conditions to perform differential expression (DE) analysis. A total of 143 differentially expressed genes (DEGs) (padj<0.05 and |L2FC|>1) were identified, including 49 upregulated and 94 downregulated genes in preterm compared to term placenta (**Table 4**). Consistent with previously studies of pregnancy complications, *NTRK2* and *BTNL9* were found to be upregulated [32], while *APLN* was downregulated in preterm versus term placentas [33] (**Figure 3**). Of all the DEGs, the most significant was *TLR7* (padj=9.87E-08), which was downregulated 2.5-fold in preterm compared to term placenta. The *TLR7* gene encodes the toll-like receptor 7 protein that detects single-stranded RNA and plays an important role in the recognition of retroviral infections and activation of innate immunity [34]. While the majority of ERVK loci examined were similarly expressed, one of the top DEGs was the ERVK11q23.3 locus (also known as ERVK-20, c11_B, and HERV-K37), which was found to be approximately four times higher in preterm compared to term placental tissue (padj=1.20E-05, L2FC=2.04) (**Figure 3**). Since TLR7 is considered essential for the control of ERVs [35], the decreased expression of TLR7 may contribute to the upregulation of the ERVK11q23.3 locus in preterm placenta. Moreover, it also supports the notion that differences in antiviral innate immune responses likely exist between term and preterm placentas [36].

**Figure 3.**
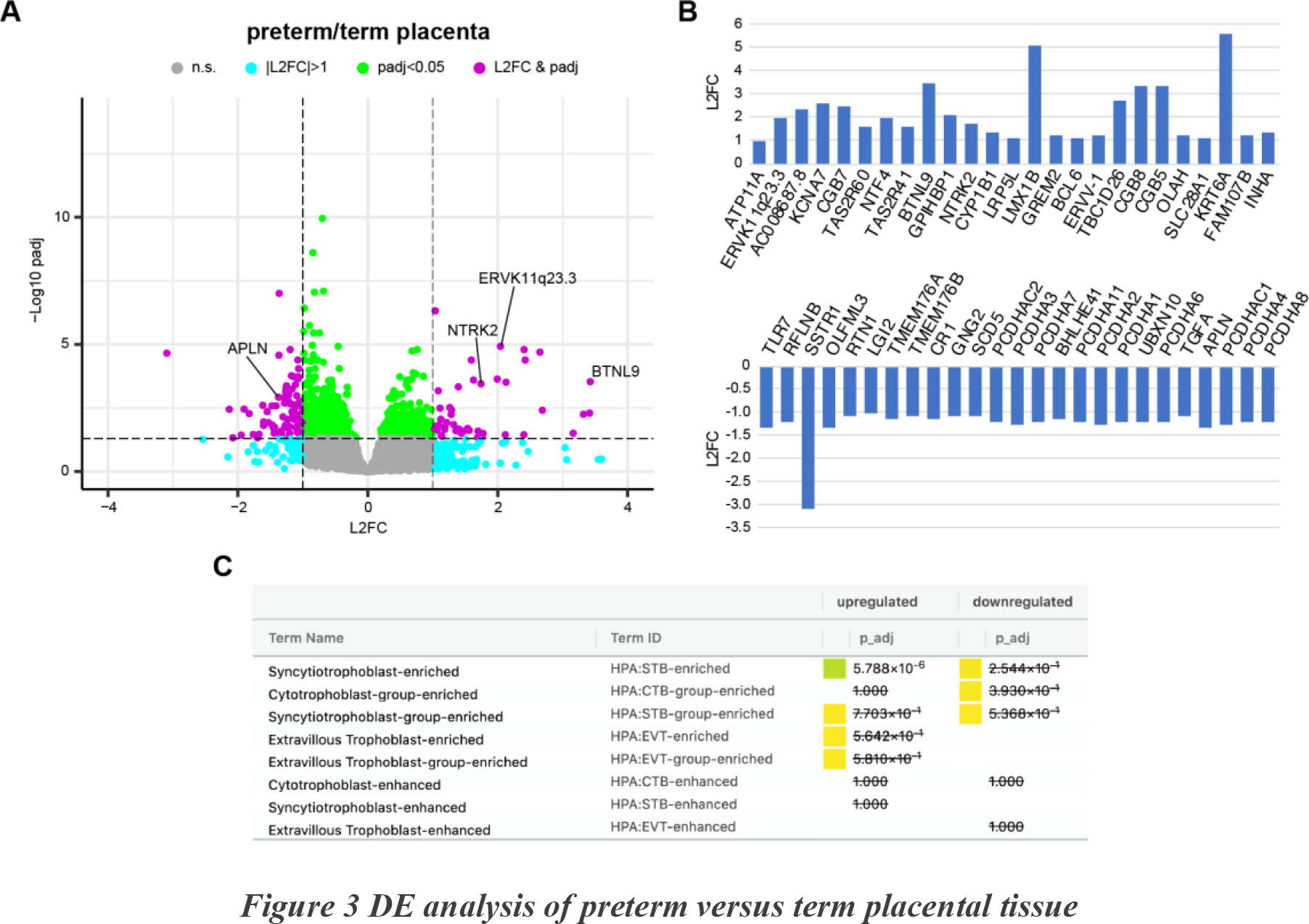
DE analysis of preterm versus term placental tissue. (A) Volcano plot of preterm versus term placenta DE results. DEGs (|L2FC| >1 and padj<0.05) are shown in purple. (B) Bar chart depicting L2FC values of the 25 most significantly upregulated (top) and downregulated (bottom) DEGs. (C) Trophoblast subtype ORA results highlighting enrichment of STB-associated genes in preterm placenta. n.s.=not significant; L2FC=Log2 fold-change; padj=adjusted p-value; STB=syncytiotrophoblast; CTB=cytotrophoblasts; EVT=extravillous trophoblasts; HPA=Human Protein Atlas.

**Table 4.**
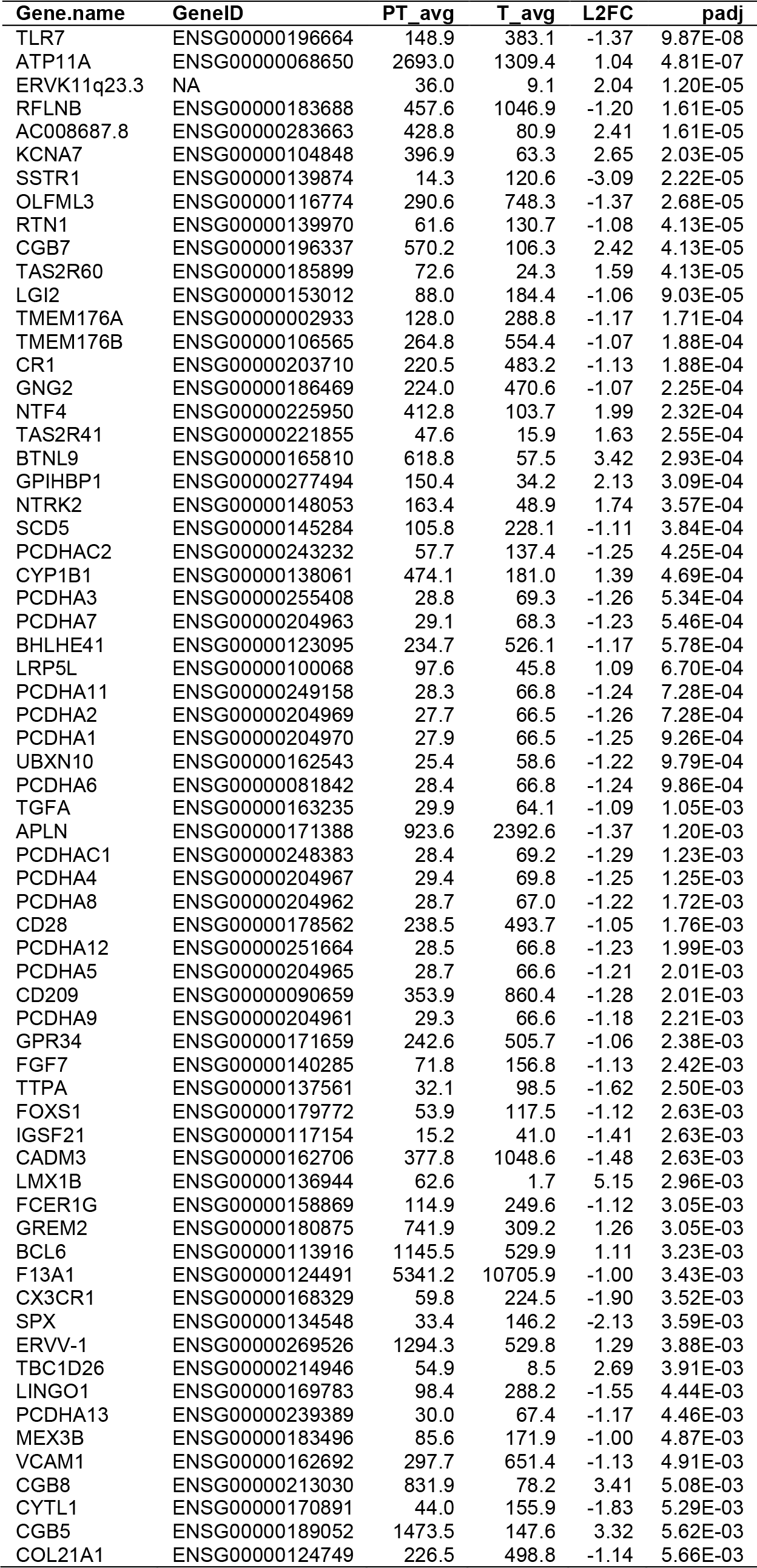
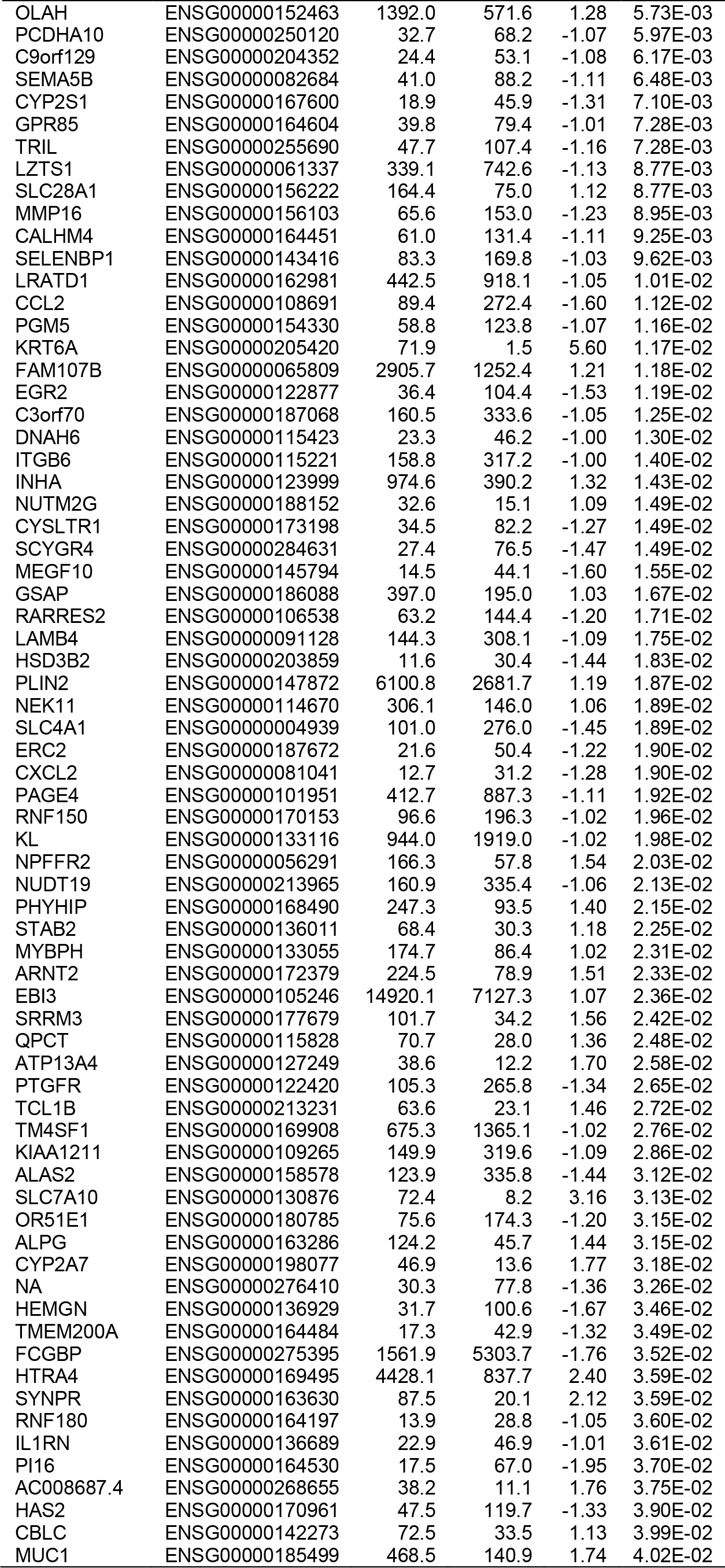
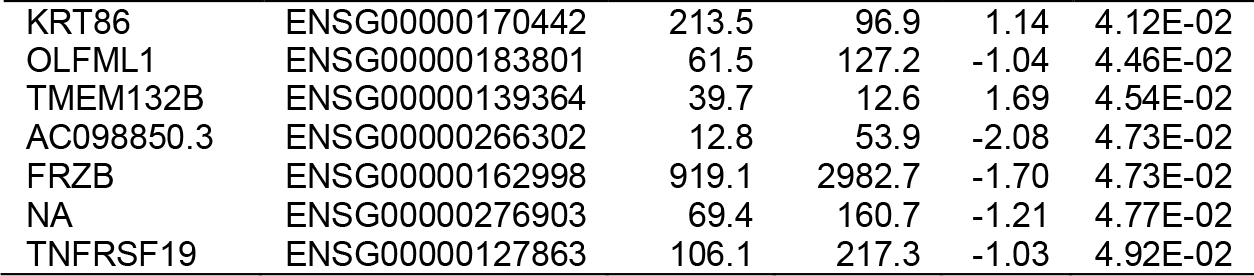
DEGs preterm/term placenta (All DEGs)

To determine whether there were differences associated with specific trophoblast subtypes between preterm and term placenta, we performed an over-representation analysis (ORA) using gene sets known to be enriched in multinucleated STBs, mononuclear CTBs, or invasive extravillous trophoblasts (EVTs). The analysis revealed that STB-associated genes (*CGB5*, *CGB7*, *CGB8*, *ERVV-1*, *INHA*, *GREM2*, *TCL1B*) were significantly over-represented in preterm-upregulated DEGs, while there was no significant enrichment of CTB or EVT associated genes found in either the upregulated or downregulated DEG sets. These results suggest that the expression differences between term and preterm placenta are predominantly associated with the STB layer rather than the other trophoblast subtypes.

### ERVK11q23.3 RNA expression is enriched in undifferentiated mononuclear CTBs

The transcriptional activity of each ERVK locus is known to vary between different tissues and cell types [30] and the placenta is a heterogeneous organ comprised of many types of cells [37]. To determine whether ERVK11q23.3 was predominantly transcribed from trophoblasts or some other placental cell type, we utilized publicly available RNA-seq data to preform DE analysis between HPTs (n=2) and bulk placenta tissue (n=5). The results showed that ERVK11q23.3 was significantly upregulated (padj=1.38E-33, L2FC=4.8) in HPTs compared to bulk placenta (**Figure 4A**, **Table 5**), indicating that placental ERVK11q23.3 RNA expression predominantly originates from trophoblast cells. An additional DE analysis using publicly available RNA-seq data from undifferentiated (n=6) and differentiated/fused (n=6) HPTs revealed that ERVK11q23.3 was significantly downregulated (padj=7.87E-21, L2FC=-2.5) in differentiated compared to undifferentiated HPTs (**Figure 4B**, **Table 6**). Collectively, these results suggest that ERVK11q23.3 RNA expression is enriched specifically within unfused mononuclear trophoblast cells of the placenta that are posed to re-populate the STB layer as it ages.

**Figure 4.**
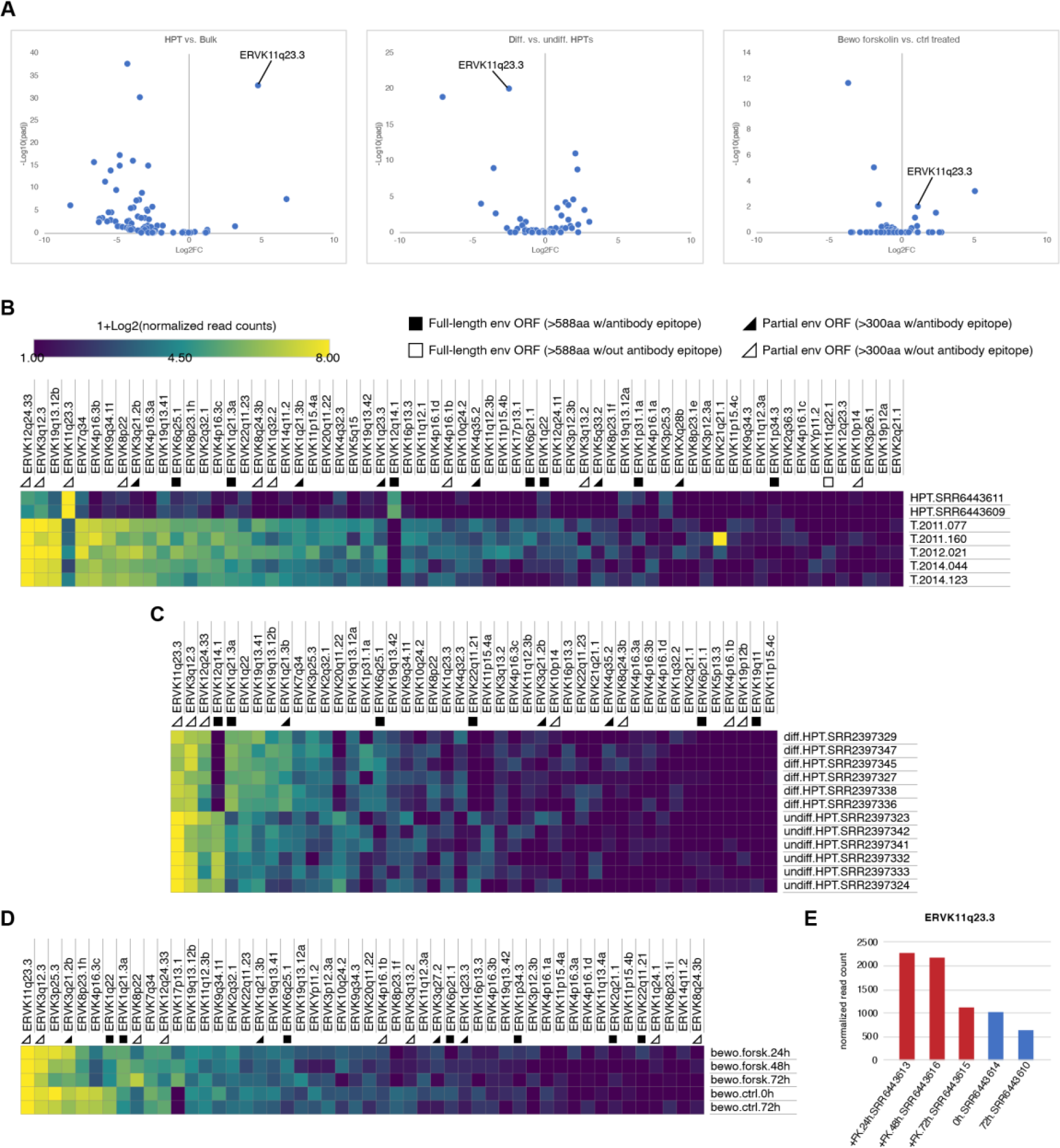
ERVK11q23.3 expression is enriched in mononuclear trophoblast cells. (A) Volcano plots of ERVK loci DE results from HPT versus bulk placenta (left), differentiated versus undifferentiated HPTs (middle), and forskolin-treated versus untreated BeWo cells. ERVK11q23.3 is significantly differentially expressed in all three analyses. (B) Heatmap depicting 1+Log2(normalized read counts) for ERVK loci with mean normalized read counts > 1 from comparison of (B) HPT versus bulk placenta, (C) differentiated versus undifferentiated HPTs, and (D) forskolin-treated versus untreated BeWo cells. (E) Bar chart of ERVK11q23.3 normalized read counts across forskolin-treated (red) and untreated (blue) BeWo samples.

**Table 5.**
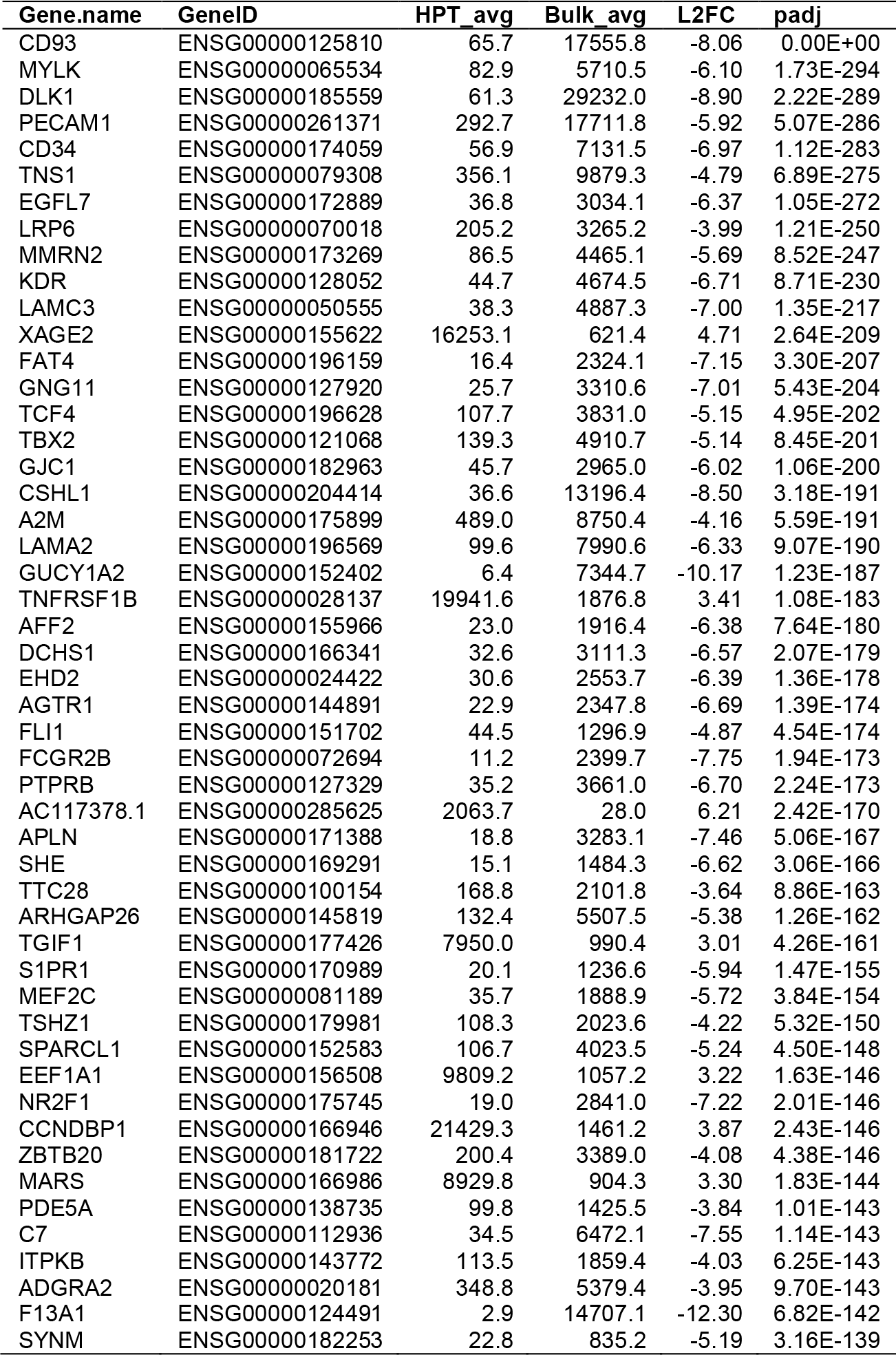
DEGs HPTs/Bulk placenta (Top 50)

**Table 6.** DEGs Diff./Undiff. HPTs (Top 50)

| Gene.name | GeneID | Diff_avg | Undiff_avg | L2FC | padj |
| --- | --- | --- | --- | --- | --- |
| ZFAT | ENSG00000066827 | 13542.5 | 3884.6 | 1.80 | 0.00E+00 |
| TGFBR3 | ENSG00000069702 | 14691.3 | 5028.2 | 1.55 | 0.00E+00 |
| CDC6 | ENSG00000094804 | 98.7 | 2046.3 | -4.38 | 0.00E+00 |
| SUMF2 | ENSG00000129103 | 2905.2 | 695.5 | 2.06 | 0.00E+00 |
| TBC1D5 | ENSG00000131374 | 1662.4 | 216.8 | 2.94 | 0.00E+00 |
| SLC30A2 | ENSG00000158014 | 6043.9 | 42.4 | 7.16 | 0.00E+00 |
| DEPP1 | ENSG00000165507 | 16767.4 | 1964.2 | 3.09 | 0.00E+00 |
| MCM7 | ENSG00000166508 | 973.4 | 4807.6 | -2.30 | 0.00E+00 |
| DHCR7 | ENSG00000172893 | 4274.8 | 184.2 | 4.54 | 0.00E+00 |
| SFN | ENSG00000175793 | 45.0 | 2899.9 | -6.02 | 0.00E+00 |
| TCL1B | ENSG00000213231 | 4919.2 | 47.3 | 6.70 | 0.00E+00 |
| LBH | ENSG00000213626 | 407.8 | 3795.1 | -3.22 | 0.00E+00 |
| LGALS13 | ENSG00000105198 | 3981.3 | 60.4 | 6.04 | 4.69E-298 |
| CDCA5 | ENSG00000146670 | 47.0 | 857.0 | -4.19 | 3.90E-293 |
| FHDC1 | ENSG00000137460 | 10601.9 | 2506.2 | 2.08 | 6.82E-293 |
| ANLN | ENSG00000011426 | 168.2 | 2003.0 | -3.57 | 1.87E-287 |
| PCNA | ENSG00000132646 | 619.4 | 3000.1 | -2.28 | 8.37E-287 |
| HSPB1 | ENSG00000106211 | 10068.8 | 2106.5 | 2.26 | 4.64E-280 |
| SDC1 | ENSG00000115884 | 33192.9 | 1471.7 | 4.50 | 5.77E-280 |
| CCNDBP1 | ENSG00000166946 | 11935.7 | 2956.6 | 2.01 | 3.16E-273 |
| DPEP2 | ENSG00000167261 | 1152.6 | 34.9 | 5.04 | 4.46E-270 |
| TTC7B | ENSG00000165914 | 2163.5 | 253.5 | 3.10 | 1.92E-263 |
| ADAMTS6 | ENSG00000049192 | 1728.9 | 155.9 | 3.47 | 5.05E-263 |
| CTSA | ENSG00000064601 | 6436.3 | 1958.8 | 1.72 | 5.73E-242 |
| SMC4 | ENSG00000113810 | 538.3 | 2024.2 | -1.91 | 1.41E-228 |
| CTNNAL1 | ENSG00000119326 | 571.6 | 2867.3 | -2.32 | 1.60E-226 |
| INHA | ENSG00000123999 | 2925.9 | 250.9 | 3.54 | 3.00E-226 |
| GCLM | ENSG00000023909 | 733.8 | 7405.7 | -3.34 | 2.75E-225 |
| ANO6 | ENSG00000177119 | 5397.0 | 17931.8 | -1.73 | 8.90E-224 |
| ARL6IP1 | ENSG00000170540 | 2479.0 | 9627.7 | -1.96 | 1.37E-220 |
| SPRR3 | ENSG00000163209 | 18.6 | 1574.5 | -6.40 | 2.39E-219 |
| PLS1 | ENSG00000120756 | 896.5 | 69.7 | 3.69 | 4.35E-218 |
| DTL | ENSG00000143476 | 28.3 | 932.6 | -5.03 | 2.24E-211 |
| WLS | ENSG00000116729 | 1376.8 | 4075.5 | -1.57 | 2.68E-210 |
| LIMA1 | ENSG00000050405 | 5421.2 | 22635.1 | -2.06 | 3.10E-210 |
| CDC25A | ENSG00000164045 | 23.9 | 725.4 | -4.90 | 4.84E-209 |
| CDT1 | ENSG00000167513 | 30.3 | 513.8 | -4.07 | 3.13E-205 |
| PSMD11 | ENSG00000108671 | 2585.1 | 5832.9 | -1.17 | 4.74E-202 |
| NDFIP2 | ENSG00000102471 | 2440.0 | 6486.5 | -1.41 | 6.81E-196 |
| CLIC3 | ENSG00000169583 | 5446.2 | 396.9 | 3.78 | 1.25E-195 |
| ENG | ENSG00000106991 | 9423.0 | 885.0 | 3.41 | 4.80E-195 |
| NAGK | ENSG00000124357 | 3637.0 | 987.8 | 1.88 | 1.13E-194 |
| NA | ENSG00000187837 | 3805.8 | 331.3 | 3.52 | 1.17E-194 |
| HOPX | ENSG00000171476 | 5546.9 | 228.6 | 4.60 | 5.99E-192 |
| SLC9A7 | ENSG00000065923 | 618.4 | 1990.4 | -1.68 | 3.19E-191 |
| ELMO1 | ENSG00000155849 | 756.1 | 26.4 | 4.84 | 3.16E-186 |
| NHLRC3 | ENSG00000188811 | 2404.6 | 270.3 | 3.15 | 1.00E-185 |
| C1orf115 | ENSG00000162817 | 19614.1 | 4048.0 | 2.28 | 3.59E-185 |
| SIDT2 | ENSG00000149577 | 5690.7 | 1022.0 | 2.48 | 8.20E-184 |
| JADE2 | ENSG00000043143 | 4314.1 | 757.3 | 2.51 | 6.11E-183 |

**Table 7.** DEGs Forskolin/Control BeWo cells (Top 50)

| Gene.name | GeneID | FSK.24h | FSK.48h | FSK.72h | FSK.avg | Ctrl.avg | L2FC | padj |
| --- | --- | --- | --- | --- | --- | --- | --- | --- |
| WNT3A | ENSG00000154342 | 19861.9 | 23783.4 | 22143.8 | 21929.7 | 454.5 | 5.59 | 1.66E-170 |
| OAF | ENSG00000184232 | 7563.0 | 8040.9 | 9505.9 | 8370.0 | 237.1 | 5.15 | 1.30E-68 |
| PLS1 | ENSG00000120756 | 748.5 | 628.2 | 585.9 | 654.2 | 6879.8 | -3.39 | 1.73E-64 |
| ST3GAL2 | ENSG00000157350 | 386.4 | 363.9 | 349.2 | 366.5 | 2237.5 | -2.61 | 9.03E-55 |
| COL9A2 | ENSG00000049089 | 151.7 | 125.3 | 127.8 | 134.9 | 1189.4 | -3.14 | 1.59E-51 |
| FIGNL2 | ENSG00000261308 | 310.5 | 230.8 | 208.3 | 249.9 | 2932.2 | -3.55 | 2.08E-48 |
| CCL28 | ENSG00000151882 | 118.3 | 113.4 | 94.7 | 108.8 | 950.4 | -3.12 | 1.65E-46 |
| FAM102B | ENSG00000162636 | 4597.3 | 5436.2 | 5794.0 | 5275.8 | 779.4 | 2.76 | 2.22E-44 |
| GPRC5A | ENSG00000013588 | 368.2 | 273.2 | 221.3 | 287.6 | 4747.0 | -4.04 | 3.64E-44 |
| AKR1B10 | ENSG00000198074 | 37.4 | 23.7 | 21.3 | 27.5 | 546.1 | -4.31 | 5.59E-42 |
| SLC15A2 | ENSG00000163406 | 55.6 | 50.3 | 41.4 | 49.1 | 736.0 | -3.90 | 6.27E-41 |
| JPH2 | ENSG00000149596 | 511.8 | 461.6 | 378.8 | 450.7 | 3073.6 | -2.77 | 8.04E-40 |
| GSDME | ENSG00000105928 | 165.9 | 150.9 | 118.4 | 145.0 | 1154.4 | -2.99 | 3.38E-39 |
| CD59 | ENSG00000085063 | 14649.6 | 15348.0 | 16166.4 | 15388.0 | 4132.1 | 1.90 | 8.99E-38 |
| PRR9 | ENSG00000203783 | 2715.9 | 7702.6 | 5388.0 | 5268.8 | 118.4 | 5.48 | 1.56E-37 |
| PRSS22 | ENSG00000005001 | 253.9 | 267.3 | 253.3 | 258.2 | 1683.0 | -2.70 | 6.95E-36 |
| COLEC12 | ENSG00000158270 | 5474.3 | 6161.1 | 6080.4 | 5905.3 | 1543.7 | 1.94 | 1.49E-35 |
| OVOL1 | ENSG00000172818 | 42941.4 | 55852.3 | 44166.9 | 47653.5 | 8023.2 | 2.57 | 2.05E-35 |
| ACHE | ENSG00000087085 | 160.8 | 109.5 | 110.1 | 126.8 | 1317.1 | -3.37 | 5.62E-35 |
| TCF7L1 | ENSG00000152284 | 56.6 | 38.5 | 33.1 | 42.8 | 800.0 | -4.22 | 1.41E-33 |
| KRT86 | ENSG00000170442 | 1760.0 | 1893.6 | 1319.8 | 1657.8 | 173.3 | 3.25 | 3.17E-33 |
| AL049629.2 | ENSG00000284969 | 7003.7 | 7682.9 | 8321.1 | 7669.2 | 1828.1 | 2.07 | 1.43E-32 |
| PLA2G15 | ENSG00000103066 | 7607.5 | 6321.9 | 6416.6 | 6782.0 | 1575.5 | 2.11 | 5.74E-32 |
| LIF | ENSG00000128342 | 112.3 | 89.7 | 86.4 | 96.1 | 683.9 | -2.83 | 5.76E-31 |
| NTN4 | ENSG00000074527 | 167.9 | 137.1 | 129.0 | 144.7 | 905.2 | -2.64 | 2.05E-30 |
| FAM110A | ENSG00000125898 | 662.5 | 598.7 | 570.5 | 610.6 | 2411.9 | -1.98 | 3.43E-30 |
| RHOV | ENSG00000104140 | 1542.5 | 1032.6 | 1239.3 | 1271.5 | 142.7 | 3.16 | 1.28E-29 |
| AC244197.3 | ENSG00000241489 | 5219.4 | 5747.9 | 6636.8 | 5868.0 | 1359.1 | 2.11 | 3.25E-29 |
| CRYBA2 | ENSG00000163499 | 551.3 | 508.9 | 454.5 | 504.9 | 2063.0 | -2.03 | 5.61E-29 |
| IDS | ENSG00000010404 | 6680.0 | 7561.6 | 8685.7 | 7642.4 | 1714.6 | 2.16 | 5.61E-29 |
| CABLES1 | ENSG00000134508 | 497.7 | 432.0 | 404.8 | 444.8 | 1945.6 | -2.13 | 6.90E-28 |
| MTMR11 | ENSG00000014914 | 251.9 | 259.4 | 224.9 | 245.4 | 1024.5 | -2.06 | 9.77E-28 |
| KCTD15 | ENSG00000153885 | 208.4 | 141.0 | 125.5 | 158.3 | 1369.1 | -3.11 | 2.83E-27 |
| SYNE3 | ENSG00000176438 | 1352.4 | 1307.8 | 1201.4 | 1287.2 | 5001.1 | -1.96 | 9.79E-27 |
| ATP8B2 | ENSG00000143515 | 821.3 | 1241.7 | 661.7 | 908.2 | 68.0 | 3.74 | 1.15E-26 |
| FGF18 | ENSG00000156427 | 398.5 | 1037.5 | 616.7 | 684.3 | 10.0 | 6.06 | 2.20E-26 |
| TP63 | ENSG00000073282 | 424.8 | 537.5 | 452.2 | 471.5 | 2002.5 | -2.09 | 3.94E-26 |
| NR4A3 | ENSG00000119508 | 1007.5 | 1402.4 | 2123.5 | 1511.1 | 103.5 | 3.87 | 5.89E-26 |
| VTN | ENSG00000109072 | 665.6 | 765.3 | 699.5 | 710.1 | 167.3 | 2.09 | 2.05E-25 |
| S100A16 | ENSG00000188643 | 242.8 | 376.7 | 325.5 | 315.0 | 1995.9 | -2.66 | 3.97E-25 |
| VLDLR | ENSG00000147852 | 1634.6 | 1455.7 | 1474.8 | 1521.7 | 5170.9 | -1.76 | 4.02E-25 |
| GPR37 | ENSG00000170775 | 638.3 | 715.0 | 642.7 | 665.3 | 2590.8 | -1.96 | 1.13E-24 |
| AL445423.3 | ENSG00000286231 | 17719.6 | 14902.2 | 13591.9 | 15404.6 | 3882.2 | 1.99 | 1.62E-24 |
| INSL4 | ENSG00000120211 | 992.3 | 1586.9 | 1558.9 | 1379.3 | 135.5 | 3.34 | 2.34E-24 |
| FAM222A | ENSG00000139438 | 1614.4 | 2083.9 | 1950.7 | 1883.0 | 440.5 | 2.10 | 2.44E-24 |
| SP6 | ENSG00000189120 | 89054.0 | 102984.1 | 70229.8 | 87422.6 | 17369.0 | 2.33 | 3.03E-24 |
| ITGB6 | ENSG00000115221 | 263.0 | 272.2 | 264.0 | 266.4 | 1109.3 | -2.06 | 3.09E-24 |
| SLC25A18 | ENSG00000182902 | 110.3 | 101.6 | 72.2 | 94.7 | 735.6 | -2.95 | 3.79E-24 |
| HSD11B2 | ENSG00000176387 | 5262.9 | 11219.6 | 5495.7 | 7326.0 | 501.2 | 3.87 | 3.79E-24 |
| THEMIS2 | ENSG00000130775 | 522.9 | 485.2 | 510.2 | 506.1 | 109.1 | 2.22 | 4.86E-24 |

**Table 8.**
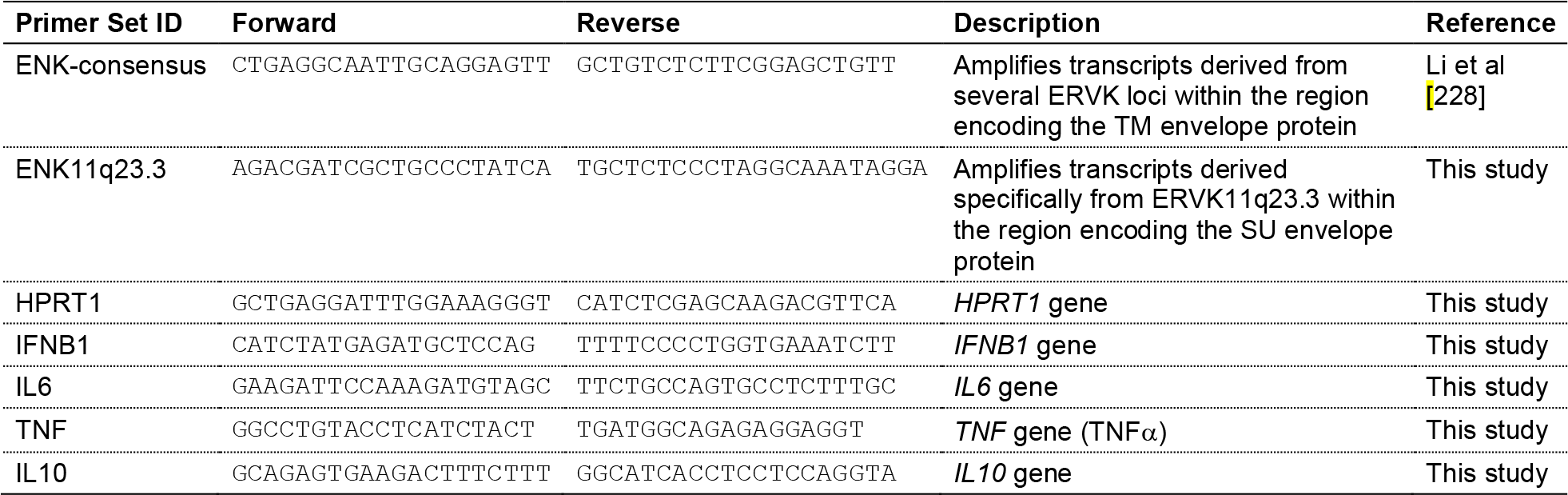
Primers.

### ERVK11q23.3 RNA expression increases in fusogenic BeWo cells

The BeWo choriocarcinoma cell line is a well-established model for studying trophoblast fusion *in vitro* following treatment with forskolin. Thus, we used publicly-available BeWo RNA-seq data from untreated (n=2) and forskolin-treated (n=3) BeWo cells [38] to investigate ERVK11q23.3 expression levels between non-fusogenic and fusogenic trophoblast cells. The DE analysis showed significant upregulation (padj=9.69E-03, L2FC=1.2) of ERVK11q23.3 in forskolin-treated fusogenic cells compared to untreated BeWo cells (**Figure 4C**, **Table 6**). Examination of the individual forskolin-treated BeWo datasets, which were derived from cells treated with forskolin for 24, 48, and 72 h [38], showed that ERVK11q23.3 expression was highest in cells at 24h and 48h, while the expression level at 72h was similar to untreated samples (**Figure 4E**). Since the fusion of BeWo cells is known to occur between 48 and 72 h after the addition of forskolin [39], these data suggest that ERVK11q23.3 may be involved in the initiation of BeWo cell fusion.

### A spliced envelope transcript is expressed from the ERVK11q23.3 locus

While full-length proviral RNA transcripts encode the *gag*, *pro,* and *pol* gene products, the envelope and accessory proteins are encoded by spliced RNA molecules [40]. Without expression of a properly spliced transcript, the envelope protein will not be produced and the resulting viruses are replication-defective [41, 42]. As a first step to assess envelope protein-coding ability, we further examined the uniquely-mapped RNA-seq reads for evidence of splicing at the ERVK11q23.3 locus. Unlike single-end sequencing, paired-end RNA-seq data can increase the alignment coverage of repetitive sequences, since uniquely-mapping mates can be used to correctly align multimapping reads. Because several regions across ERVK11q23.3 are non-unique and have low mappability (**Figure 5**), we relied on the use of paired-end RNA-seq data to identify spliced ERVK11q23.3 transcripts. First, we manually examined the unmapped mates of reads uniquely mapping to the ERVK11q23.3 locus from publicly-available BeWo paired-end RNA-seq data (n=2) [43]. In total, we uncovered the presence of at least four splice sites and three distinct spliced transcripts generated from the ERVK11q23.3 locus (**Figure 5**). This included a (1) full-length transcript with a 437aa truncated *gag ORF*, (2) single-spliced transcript with a 438aa truncated *env ORF*, (3) single-spliced transcript with 44aa truncated *Np9* (an ERVK accessory protein) ORF, and (4) double-spliced transcript with a 74aa full-length *Np9* ORF (**Figure 5**). Taken together, these results suggest that the ERVK11q23.3 locus has the ability to encode a truncated envelope protein.

**Figure 5.**
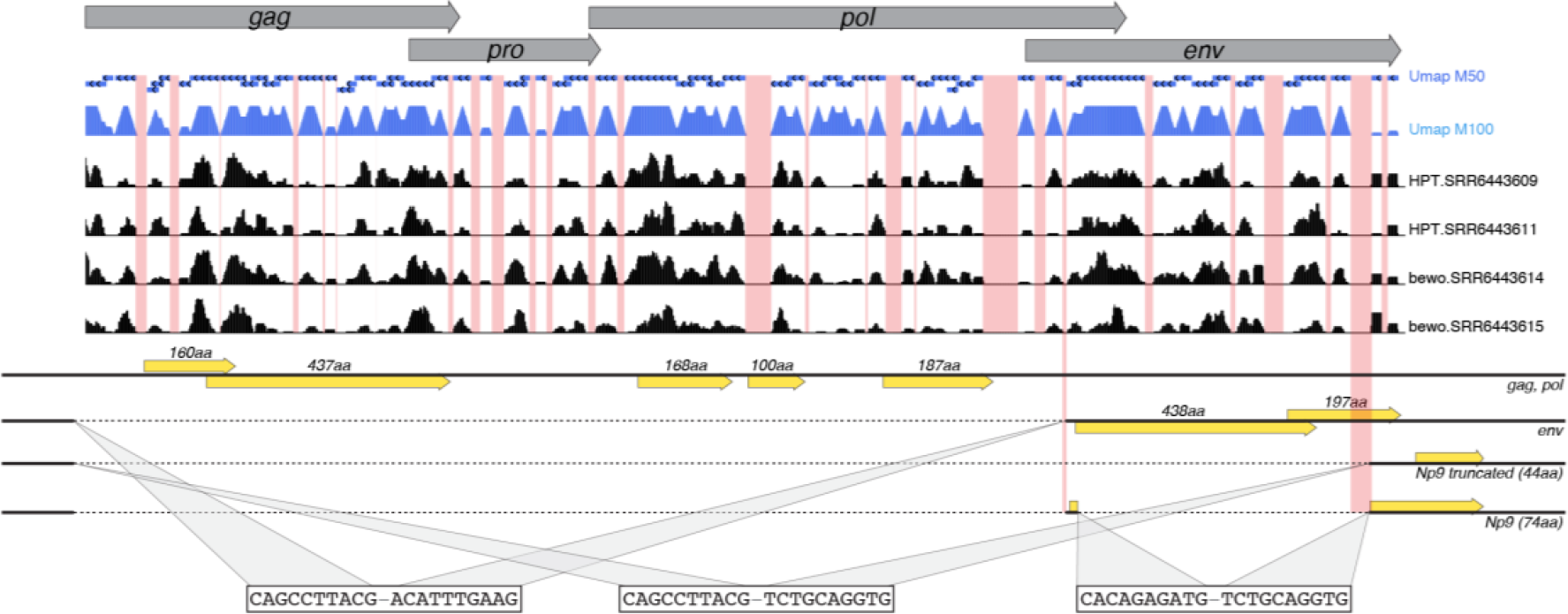
Representation of ERVK11q23.3 spliced transcripts. Schematic of the ERVK11q23.3 locus, including proviral genes (grey). Umap tracks (blue) highlight the regions with low mappability (red). Tracks of mapped RNA-seq data from HPT (n=2) and BeWo (n=2) samples are included (black). Unmapped mate reads from BeWo RNA-seq data were analyzed and identified the presence of at least four splice sites and three distinct spliced transcripts generated from the ERVK11q23.3 locus. Putative ORFs for each transcript are shown in yellow.

### The predicted ERVK11q23.3 envelope protein contains FP and ISU domains, but lacks a membrane spanning region

The retroviral envelope gene product encodes a polyprotein that is cleaved by cellular proteases to yield mature surface unit (SU) and transmembrane (TM) envelope proteins [44], with the FP and ISU domains located on the TM protein [6, 7]. Similar to exogenous retroviruses, the ERVK ancestral-predicted TM protein contains functional FP and ISU domains [18–21], but the presence of these domains within the putative ERVK11q23.3 envelope protein is currently unknown. Therefore, we examined the amino acid sequence of two overlapping ORFs identified within the ERVK11q23.3 single-spliced envelope transcript for the presence of the FP and ISU domain sequences. The first and longest ORF is predicted to encode a partial 438aa polyprotein containing the SU and part of the TM envelope protein, whereas the second ORF is 197aa long and corresponds to the remainder of the TM subunit. While the 438aa ORF protein sequence contains a Furin cleavage site [31], FP [45], and ISU domain [21], the 197aa ORF protein sequence contains the membrane spanning region (MSR) and anti-ENK epitope (**Figure 6A**). A hydrophobicity plot along the merged protein sequences confirmed the presence of both a FP and MSR, which are known to be hydrophobic [6, 7] (**Figure 6B**). When compared to the ancestral-predicted ERVK-env sequence [28], a 2bp deletion at the end of the ISU domain appeared to cause a frameshift and subsequent premature stop codon in the ERVK11q23.3 envelope protein. Notably, a +1 frameshift near the end of the first ORF could result in translation of full-length envelope protein from this locus (**Figure 6C**). However, Sanger sequencing of cDNA from HPTs (n=6 clones) showed no insertions, deletions or splicing events that would allow a full-length envelope protein to be translated (**Figure 6C**). This suggests that the single-spliced transcript from the ERVK11q23.3 locus encodes a 438aa truncated envelope protein that may be secreted due to the lack of a MSR. Nonetheless, the presence of intact FP and ISU domains indicates that this protein may still be involved in trophoblast fusion and/or immunosuppression.

**Figure 6.**
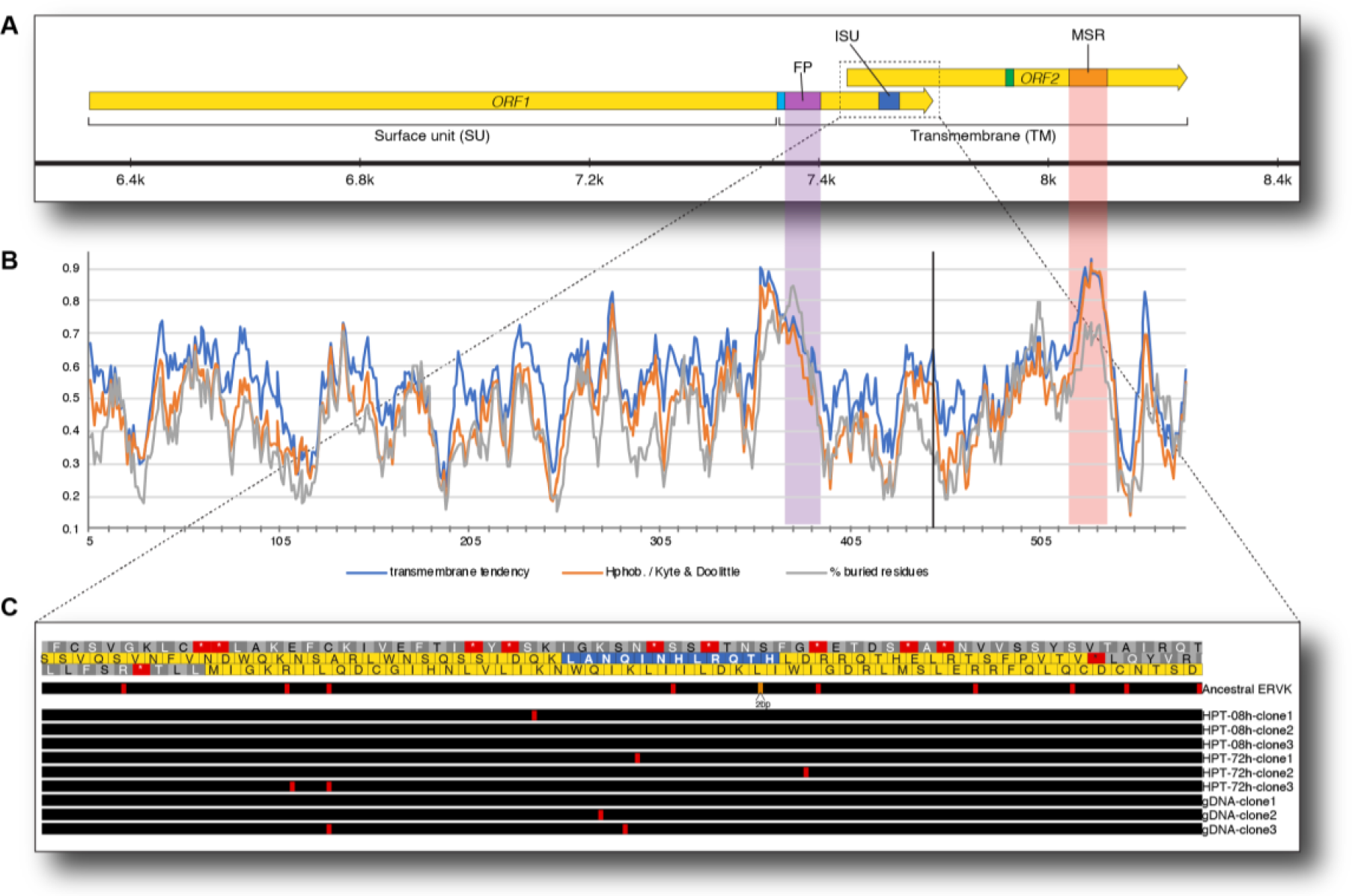
Predicted ERVK11q23.3 envelope protein contains FP and ISU domains, but lacks a MSR. (A) Schematic of the two overlapping ORFs (yellow) in the ERVK11q23.3 envelope gene. ORF1 (left), contains a well-documented Furin cleavage site (cyan), FP (purple), and ISU domain (blue), while ORF2 (right), contains the ERVK-env monoclonal antibody epitope (green), and MSR (orange). (B) Plot of hydrophobicity (orange), transmembrane tendency (blue), and % buried residues (gray), across 9aa windows of the predicted envelope protein. (C) Comparison to the ancestral predicted ERVK sequence (top track) revealed a 2bp deletion in ERVK11q23.3 envelope gene (orange). Examination of cDNA clones from HPTs showed several mismatched bases (red), but no insertions, deletions or splicing events that would result in translation of a full-length envelope protein.

### Knockdown of ERVK11q23.3 envelope has no effect on HPT and BeWo cell fusion levels

Since certain truncated ERV envelope proteins have previously been shown to influence trophoblast fusion, while others do not have this ability [46–48], we sought to elucidate the effect of ERVK11q23.3 expression on trophoblast cell-cell fusion. For this, small interfering RNAs (siRNAs) targeting the envelope gene were used to knockdown (KD) ERVK11q23.3 expression in both fusogenic BeWo and HPT cells. Greater than 70% of both BeWo and HPT cells showed successful uptake of a fluorescently labeled siRNA transfection control (**Figure 7A**). An siRNA targeting no known sequence in the human transcriptome (siNC1) was included as a negative control, and an siRNA targeting the HPRT1 gene (siHPRT1) was used as a positive control. When transfected with siHPRT1 both BeWo (n=4, p=3.38E-06) and HPT (n=3, p=2.25E-02) showed a ∼82% reduction in HPRT1 RNA level compared to siNC1 transfected cells (**Figure 7B**). Out of three different siRNAs tested (siENK13.13, siENK13.7, and siENK13.34), siENK13.34 was determined to be the most effective, reducing ERVK11q23.3 expression by ∼70% (n=4, p=1.21E-04) compared to siNC1 transfected cells (**Figure 7C**). Thus, siENK13.34 was utilized in all subsequent experiments (referred to as siENK hereinafter). To assess the effect of ERVK11q23.3 silencing on cell-cell fusion, IF staining of the plasma membrane marker, E-cadherin (CDH1), was used to calculate the percent fusion for both siENK (n=4) and siNC1 (n=4) transfected cells. Transfection of siENK significantly reduced ERVK11q23.3 expression by ∼74% in BeWo cells (p=1.01E-04) and by ∼60% in HPTs (p=2.18E-02). However, this decrease did not appear to be associated with any significant changes in cell fusion levels in either BeWo or HPT cells (**Figure 8**). Despite the role of other ERVs in trophoblast fusion [48], these results suggest that ERVK11q23.3 envelope expression is not involved in the cell-cell fusion normally observed in forskolin-treated BeWo cells or primary HPTs.

**Figure 7.**
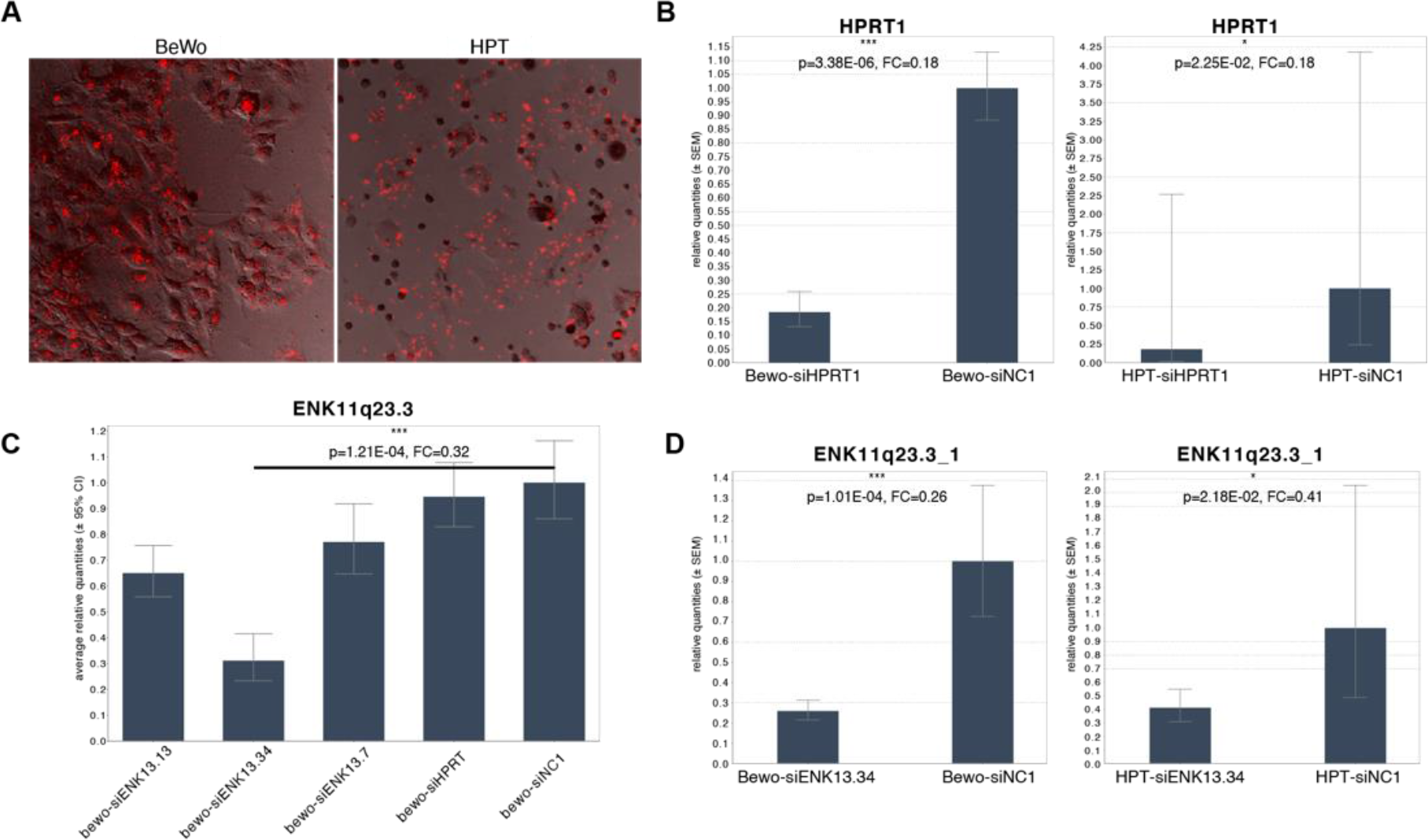
Validation and optimization of ERVK11q23.3 siRNA knockdown. (A) Microphotograph of cells transfected with the fluorescently-labeled siRNA control. (B) HPRT1 relative quantities determined via qRT-PCR from cells transfected with siHPRT1 or siNC1. (C) ERVK11q23.3 envelope transcript (ENK11q23.3) relative quantities determined via qRT-PCR from BeWo cells transfected with either siENK13.13, siENK13.34, siENK13.7, siHPRT1, or siNC1. (D) ENK11q23.3 relative quantities determined via qRT-PCR from cells transfected with siENK13.34 or siNC1. Samples were normalized to GAPDH; error bars reflect SEM.

**Figure 8.**
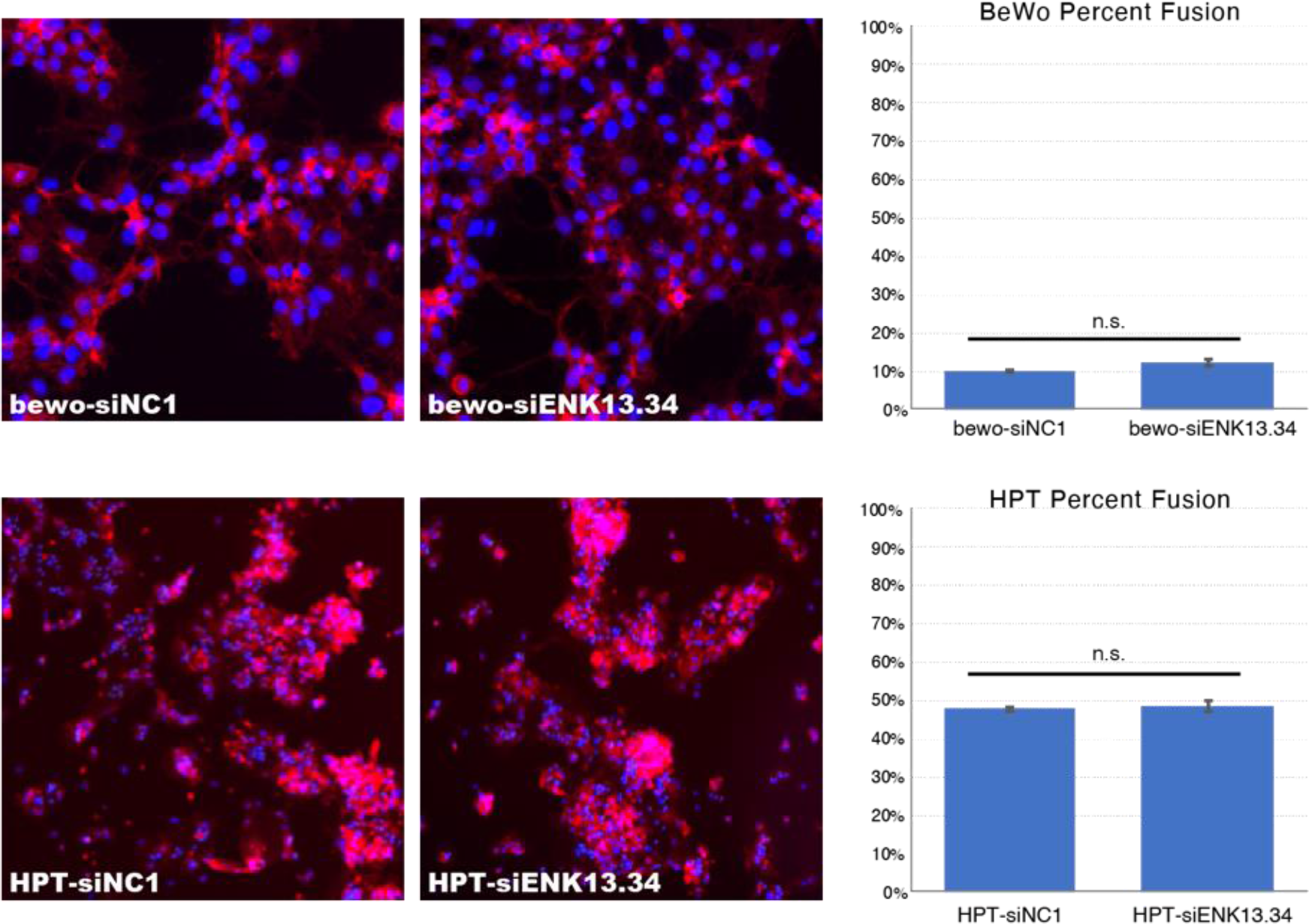
siRNA KD of ERVK11q23.3 showed no change in trophoblast fusion levels. Representative CDH1 (red) and DAPI (blue) immunostained microphotographs (n=5 per well) from BeWo (top) and HPT (bottom) cells transfected with siNC1 (n=3) (left) and siENK (n=3) (right). Bar charts depict fusion percentage (± standard deviation) calculated for each condition.

### Knockdown of ERVK11q23.3 envelope decreased expression of type I interferon and antiviral immune response in trophoblast cells

Innate immune responses can be divided into two categories: acute inflammatory responses and antiviral responses. The inflammatory response is marked by an induction of small signaling molecules called cytokines, including interleukin-6 (IL6) and tumor necrosis factor α (TNFα) [49, 50], while the antiviral response is characterized by the release of type I interferons (IFN), including IFNB1 [51]. To assess the putative role of ERVK11q23.3 in the innate immune response, we compared the expression level of several immune modulatory genes between siENK (n=3) and siNC1 transfected (n=3) cells via qRT-PCR. Additionally, since siRNA alone is capable of inducing double-stranded RNA (dsRNA) triggered innate immune responses [52], we also compared siNC1 (n=3) to mock (n=3) transfected cell to assess the effect of siRNA transfection on gene expression levels. Indeed, compared to mock transfected cells, HPTs transfected with siNC1 showed significant upregulation of antiviral type I IFN, IFNB1 (12-fold, p=2.62E-2 (**Figure 9**), suggesting that siRNA transfection likely induces a dsRNA triggered antiviral response. However, when HPTs were transfected with siENK (n=3), a 78% reduction in IFNB1 expression (0.22- fold, p=8.66E-04) was observed compared to siNC1 (n=3) (**Figure 9**). This suggests that the loss of ERVK11q23.3 diminishes the antiviral cytokine expression normally initiated by siRNA transfection. To a lesser extent, siRNA transfection of HPTs also induced expression of the pro-inflammatory cytokines, IL6 (3.46-fold, p=4.34E-03) and TNFα (3.51-fold, p=3.14E-02) (**Figure 9**). Unlike IFNB1, IL6 and TNFα expression levels were not significantly different between siENK and siNC1 transfected HPT cells (**Figure 9**), suggesting that loss of ERVK11q23.3 expression has no significant effect on the pro-inflammatory response elicited by siRNA transfection in HPTs.

**Figure 9.**
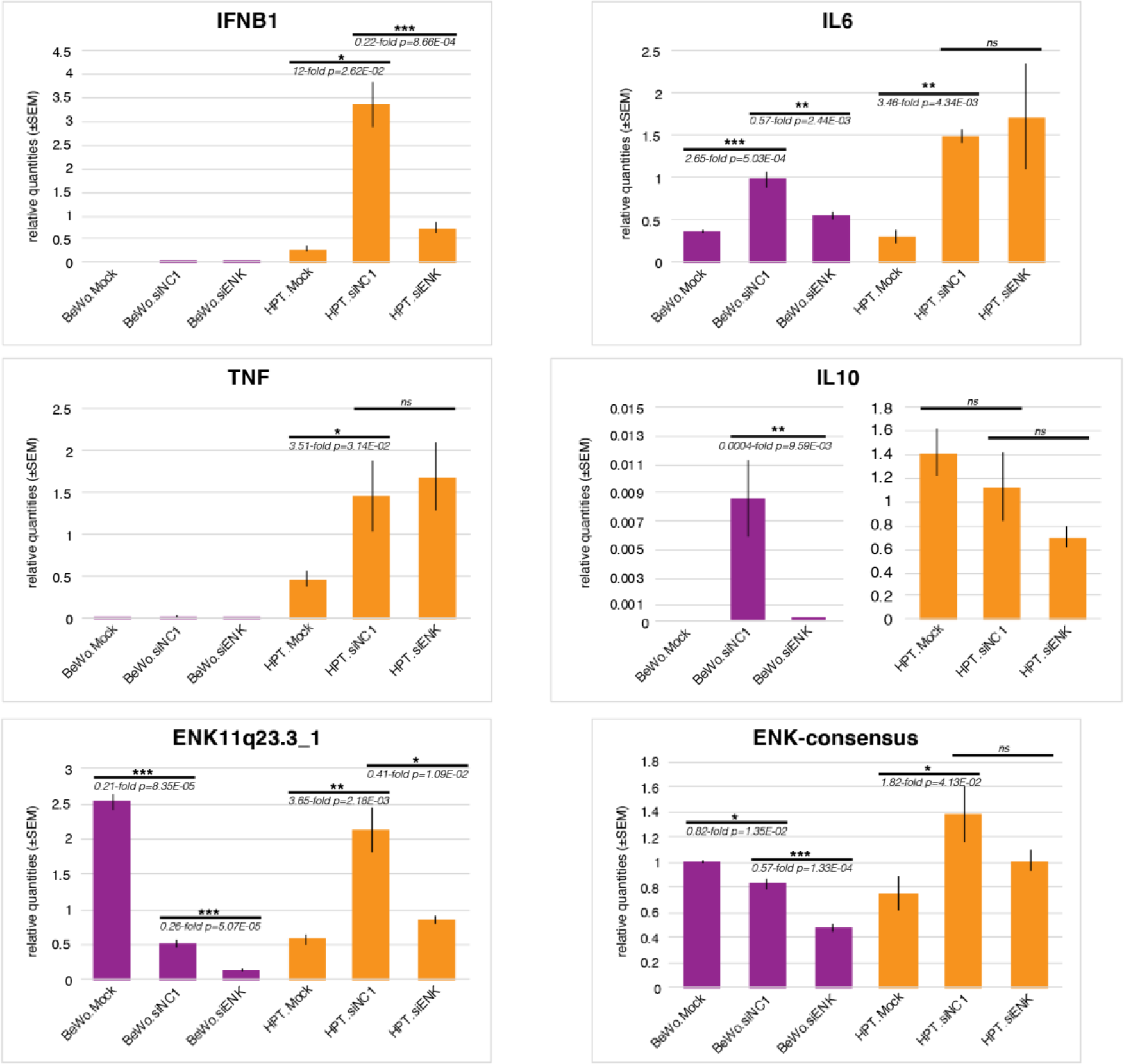
siRNA KD of ERVK11q23.3 decreased expression of key immunomodulatory genes. Relative gene expression levels determined via qRT-PCR of several well-known immunomodulatory genes from BeWo (purple) and HPT (orange) cells transfected with siENK13.34 (light colored, n=3) and siNC1 (dark colored, n=3). Samples were normalized to GAPDH and scaled to siNC1 negative control samples. Error bars reflect SEM.

Similar experiments were conducted with BeWo cells, but IFNB1 and TNFα expression levels were undetectable in all samples (**Figure 9**), indicating that siRNA transfection failed to induce an IFNB1- mediated antiviral and/or a proinflammatory cytokine response. However, there was a significant increase in IL6 (2.65-fold, 5.03E-04) and IL10 expression levels in siNC1 (n=3) compared to mock transfected (n=3) BeWo cells (**Figure 9**). While IL6 is often considered a proinflammatory cytokine, in the absence of TNFα and other stress agents, it can induce the expression of immunosuppressive factors, including IL10 [53–55]. Thus, these results suggest that siRNA transfection of BeWo cells increased the expression of anti-inflammatory cytokines which are known to promote immunosuppression. However, KD of ERVK11q23.3 in BeWo cells significantly reduced levels of IL6 (0.57-fold, p=2.44E-03) and IL10 (0.0004-fold, p=9.59E-2 compared to siNC1 transfected cells (**Figure 9**), indicating that ERVK11q23.3 expression facilitates upregulation of these anti-inflammatory cytokines. Collectively, these results show that ERVK11q23.3 expression mediates the upregulation of antiviral and anti-inflammatory cytokines induced via siRNA transfection of HPTs and BeWo cells, respectively. Thus, altered expression levels of ERVK11q23.3 in trophoblast cells at the maternal-fetal interface may result in aberrant antiviral and/or anti-inflammatory maternal immune responses during pregnancy.

## Discussion

It is well-recognized that ERV viral-like proteins can be co-opted to play important biological roles during normal human placentation. While it is currently known that ERVK-env protein is expressed in the placenta and that the ancestral-predicted envelope protein possesses fusogenic and immunosuppressive functions [11,18–21], the characteristics and putative function of natively expressed ERVK-env protein during human placentation remains unclear. To assess the putative fusogenic and/or immunosuppressive role of the ERVK-env in the placenta, a more thorough characterization of placental ERVK expression including, locus-specific transcription levels, splicing, and envelope protein-coding potential is required. In this study, we comprehensively examined locus-specific ERVK transcription across several different human placental tissues and cell types. Through a combination of RNA-seq and siRNA KD analyses, we identified the expression of a single ERVK locus, ERVK11q23.3, as (1) being significantly upregulated in preterm placenta, (2) predominantly expressed by mononuclear trophoblasts, (3) capable of encoding a truncated viral-like envelope protein, and (4) contributing to the expression cytokines involved in both antiviral and anti-inflammatory innate immune responses in HPTs and BeWo cells, respectively.

While abnormal placental ERVW and ERVFRD expression has been linked to several pregnancy complications [56–60], an association between aberrant placental ERVK expression and pathological pregnancy conditions has not yet been shown. To our knowledge, this is the first study to evaluate ERVK locus-specific expression levels in human placental samples, and specifically identify that the ERVK11q23.3 locus is significantly upregulated in preterm compared to term placental tissue. This discovery was only possible via the implementation of a locus-specific strategy that examined transcription of each ERVK proviral insertion, since numerous ERVK loci were found to be similarly expressed between term and preterm placenta. Previous reports using similar locus-specific strategies have shown that ERVK11q23.3 is transcribed within germ cell tumors [30], HIV-1 infected HeLa cells [61], as well as embryonic and induced pluripotent stem cells [62]. Since ERVK11q23.3 is predominantly expressed by mononuclear CTBs, it is possible that this cell type is more abundant in preterm placental tissue. However, the DEGs identified between preterm and term placenta and the subsequent ORA results suggest that CTB abundance was equal across the two groups.

Previous studies have demonstrated that ERVK-derived transcripts and viral-like proteins are expressed during human placentation [11, 63]. These reports are consistent with our RNA-seq and immunostaining results, which highlight placental ERVK RNA and ERVK-env protein expression, respectively. However, some notable differences were observed between our ERVK-env IHC staining results and those reported by a previous study using the same monoclonal antibody [11]. While Kammerer et al. showed ERVK-env staining predominantly localized to mononuclear CTBs within placental tissue, we observed staining primarily localized to the multinucleated STB layer. Since we followed the IHC procedure described by Kammerer et al., we speculate that the different staining patterns observed were due to lack of CTBs in the tissue sections examined or possibly variability between the antibody lots used. Nonetheless, our IF staining results showed strong ERVK-env membrane staining of mononuclear HPTs, which is consistent with the CTB expression previously described [11]. Notably, the envelope ORF of the ERVK11q23.3 locus is not predicted to contain the antibody epitope, suggesting that the protein staining detected via IHC and IF is not derived from the ERVK11q23.3 locus. A total of twelve ERVK loci expressed in bulk placenta are predicted to possess an envelope ORF containing the ERVK antibody epitope. Thus, the ERVK-env protein detected via the monoclonal antibody (HERM-1811-5) used in this study is likely derived from one or more of these loci and may explain the variability in expression between preterm placentas.

It is well established that full-length exogenous retroviral RNA transcripts encode the *gag* and *pol* gene products, while the envelope and accessory proteins are encoded by spliced RNA molecules [40]. Additionally, similar splicing has been documented for several ERVK proviruses as well [64–66]. This is consistent with the results from our splicing analysis, which revealed the expression of several spliced transcripts from the ERVK11q23.3 locus, including a single-spliced envelope transcript. Previous studies have shown that without expression of a properly spliced proviral transcript, the envelope protein cannot be encoded [41, 42]. While expression of a single-spliced envelope transcript is consistent with envelope protein expression, confirmation that a truncated envelope protein is still encoded from the ERVK11q23.3 locus is needed using an alternative ERVK-env antibody or another protein-based approach. Notably, we identified a 2bp deletion relative to ancestral-predicted ERVK that would have introduced a premature stop codon within the envelope ORF, indicating that the putative ERVK11q23.3 envelope protein is truncated. However, the presence of a secondary overlapping ORF suggests expression of a full-length envelope protein from this locus is still possible via a ribosomal frameshift event, which are well-documented in retroviruses [67].

Even though placental ERVK expression has been hypothesized to aid in maternal immunomodulation [11], our study implicates its expression in trophoblast immune regulation. While the mechanism remains unclear, expression of ERVK11q23.3 viral-like RNA and/or protein sequences was able to modulate the expression levels of cytokines associated with both antiviral and immunosuppressive immune responses. Similar to exogenous retroviral infections, previous studies have shown that ERVs can induce innate antiviral immune responses when expressed in certain tissues and cell lines [68–70]. These reports are consistent with our findings that the loss of ERVK11q23.3 in HPTs significantly reduced expression of *IFNB1*, a type I IFN involved in the antiviral immune response. Activation of type I IFNs has been shown to exacerbate systemic and uterine proinflammatory cytokine production and increase susceptibility to inflammation-induced preterm birth in mice [71]. Since the loss of ERVK11q23.3 expression significantly reduced *IFNB1* expression in HPTs, upregulation of placental ERVK11q23.3 expression may enhance the type I IFN response and reduce the inflammatory challenge required for the induction of preterm birth. This is further supported by our DEG results showing that ERVK11q23.3 is significantly upregulated in preterm placental tissue.

In addition to eliciting an antiviral immune response, retroviral infections are frequently accompanied by immunosuppression, which allows retroviruses to escape host immunologic defenses. This immunosuppression is largely attributed to the retroviral TM envelope protein, which contains peptide sequences that functionally suppress immune effector cells [7]. Several ERV-derived envelope protein sequences, including the ancestral-predicted ERVK-env sequence, are known to impair immune responsiveness in a similar manner to retroviral TM envelope proteins [10,21,47,72,73]. Thus, placental ERVK expression may also facilitate maternal immunosuppression required throughout normal pregnancy as mentioned above [11]. Our findings that siRNA KD of ERVK11q23.3 in BeWo cells significantly reduced the levels of the immunosuppressive cytokines, IL10 and IL6, further supports this notion. However, these same results were not observed with siRNA KD of ERVK11q23.3 expression in HPTs, which predominantly affected antiviral cytokine expression. These different cytokine responses suggest that BeWo choriocarcinoma cells might not accurately recapitulate ERVK11q23.3 function in normal trophoblast cells. It also suggests that ERVK11q23.3 expression may have multiple functions and/or its function is context dependent. For instance, ERVK11q23.3 RNA expression could elicit an antiviral immune response, while the ERVK11q23.3 envelope protein induces an immunosuppressive response. Another possibility is that ERVK11q23.3 only elicits an antiviral immune response in the presence of adequate IFNB1 and TNFα, which are expressed at much higher levels in HPTs compared to BeWo cells. To help clarify this, examination of additional cytokines and the quantification of both ERVK11q23.3 RNA and protein from HPTs and BeWo cells should be performed.

Besides ERVK11q23.3, our locus-specific ERVK analysis revealed high expression levels of a number of other loci within placental tissues and cells. Notably, two of the most highly expressed loci in bulk placenta, ERVK12q24.33 and ERVK19q13.12b, are located antisense within the introns of placentally expressed ZNF genes, *ZNF140* and *ZNF420*, respectively. Thus, expression of ERVK12q24.33 and ERVK19q13.12b can produce antisense transcripts to unspliced ZNF transcripts, which may inhibit translation of associated ZNF proteins through complementary binding and induction of the RNA interference pathway [74]. Several ERVK loci highly expressed in HPTs were not detected within bulk placental samples, including ERVK12q14.1, which was shown to be significantly upregulated in HPTs compared to bulk placenta and in undifferentiated compared to differentiated HPTs. Unlike ERVK11q23.3, ERVK12q14.1 contains a full-length envelope ORF and is predicted to encode an ERVK-env protein possessing a MSR and the antibody epitope. Based on these characteristics, we suspect that the ERVK-env IF staining observed along the membrane of mononuclear HPTs is likely derived from this locus and may facilitate trophoblast fusion. Since ERVK12q14.1 expression was not detected within any of the BeWo samples examined, its putative role is likely enhanced in or specific to HPTs. This was the case for cell fusion, which we show occurs much more in HPTs compared to forskolin treated BeWo cells. To assess its putative fusogenic role, a similar siRNA KD approach specifically targeting the ERVK12q14.1 locus within HPT cells and/or a transgene overexpression approach in BeWo cells should be used.

In conclusion, we showed that ERVK11q23.3 transcription is upregulated in preterm placenta and facilitates antiviral cytokine gene expression in trophoblast cells, suggesting that expression of this element helps activate an antiviral immune response in the placenta. Moreover, our analysis revealed that ERVK11q23.3 has the ability to encode a partial envelope protein, however, it is still unclear whether this protein is generated and if it is involved in the innate immune response we observed in siRNA transfected trophoblast cells. Aberrant ERVK11q23.3 expression levels may contribute to pregnancy complications and/or diseases affecting innate antiviral immune response and should be further investigated. Thus, the specific mechanism underlying ERVK11q23.3-mediated antiviral cytokine expression, including the role of ERVK11q23.3 RNA and proteins, should be further interrogated in future investigation.

## Methods

### Placental tissues and cells

Deidentified human placental samples were collected by and acquired through the Labor and Delivery Unit at the Oregon Health and Science University Hospital and deposited into a repository under a protocol approved by the Institutional Review Board with informed consent from the patients. A total of five different term placenta samples collected from healthy cesarean section term births without labor (ranging from 38.9 to 41.3 gestational wks), and five preterm placental samples (ranging from 33.3 to 36.4 gestational wks) were used for RNA isolation and RNA-seq based analyses. Further, banked formalin-fixed paraffin-embedded tissues (FFPE) tissues collected from human term (n=4) and preterm (n=4) placentas were used for IHC staining. Frozen vials of HPTs consisting of highly purified CTB cells used for the IF and siRNA KD experiments were isolated from human term placental tissue using a Percoll gradient as previously described [25].

### IHC staining

Paraffin sections were deparaffinized and rehydrated through xylene and graded alcohol series, then washed for 5 min in running tap water. Antigen unmasking was performed using sodium citrate (pH 6.0) buffer in a pressure cooker for 20 min, followed by washing with three changes of PBS. An endogenous enzyme block was performed by incubating sections in 0.3% hydrogen peroxide for 10 min and similarly washed. Nonspecific protein binding was blocked by incubating sections in 5% horse serum for 30 min. The mouse monoclonal antibody specific to the ERVK-env TM protein (Austral Biologicals, HERM-1811-5) was diluted 1:250, and the mouse IgG2A isotype control was diluted to an equivalent concentration. Tissue sections were incubated in primary antibody dilutions for 2 h at room temperature (RT). Mouse IgG H+L (Vector Labs, BA-2000) and biotinylated secondary antibody dilutions were prepared at 1:250 in PBS + 1% BSA. The sections were incubated in secondary antibody dilution for 1 h at RT, then washed in three changes of PBS. VECTASTAIN Elite ABC HRP Kit (Vector Laboratories, PK-6100) and ImmPACT DAB Peroxidase HRP substrate (Vector Labs, SK-4105) were used according to the manufacturer’s instructions. All tissue sections were stained in a single batch using the same incubation times. Nuclei were counterstained with hematoxylin (Electron Microscopy Sciences, 26043-05), and imaged using a brightfield microscope.

### Transcriptional analysis of human ERVK loci

The UCSC table browser RepeatMasker track was used to extract a bed file of all genomic regions identified as HERVK-int, LTR5A, LTR5B, or LTR5_Hs (filtered by repNames: HERVK-int* OR LTR5A* OR LTR5B* OR LTR5_Hs*). Nearby regions (within 2000bp) were merged into single loci using Bedtools merge (options: -s -d 2000 -c 4,5,6 -o collapse,sum,distinct -delim “;”). Solo LTR elements were discarded, and only loci containing at least one “HERVK-int” annotation were examined within the subsequent analysis. RNA-seq raw fastq files were trimmed of low-quality and adapter sequences using Trimmomatic [75] and mapped to the human (GRCh38) reference genome using Bowtie2 [76] with --very-sensitive parameter. Resulting BAM files were filtered to remove low quality and multi-mapped reads (MAPQ ≥10) using samtools [77] view -q 10. A custom gtf file including the 124 ERVK loci and ENSEMBL human genome protein-coding gene annotations (Homo_sapian.GRCh38.98.protein_coding) was used to generate raw read count tables (n=20128 genes) using featureCounts [78] (–primary). DEseq2 [79] was used to generate VST-normalized read counts for transcriptomic comparison. The default settings of DEseq2 were used for DE analyses and genes with padj<0.05 & Log2FC>|1| were identified as DEGs. A total of four separate DE analyses were carried out, including (1) preterm (n=5) vs. term (n=5) human placenta samples, (2) HPTs (n=2) vs. bulk term placenta (n=5), (3) differentiated (n=6) vs. undifferentiated (n=6) HPTs, and (4) untreated (n=2) vs. forskolin-treated (n=3) BeWo cells. The human term and preterm RNA-seq data was generated in-house, while all other RNA-seq data was publicly available and downloaded from NCBI SRA using the SRA Toolkit (http://ncbi.github.io/sra-tools/). The following publicly available RNA-seq data was used in DE analyses described above: Bulk preterm placenta (SRR13632931, SRR13632932, SRR13632933, SRR13632934, SRR13632935); Bulk term placenta (SRR12363244, SRR12363245, SRR12363246, SRR12363247, SRR12363248); HPTs (SRR6443609, SRR6443611); Undiff. HPTs (SRR2397323, SRR2397324, SRR2397332, SRR2397333, SRR2397341, SRR2397342); Diff. HPTs (SRR2397327, SRR2397329, SRR2397336, SRR2397338, SRR2397345, SRR2397347); BeWo (SRR6443614, SRR6443610); Forskolin-treated BeWo (SRR6443613, SRR6443615, SRR6443616); paired-end BeWo (SRR9118949, SRR9118950).

### Splicing and ORF analysis

Paired-end BeWo RNA-seq data was used for further assessment of the *ERVK11q23.3* locus. Unmapped mates of reads uniquely mapping to ERVK11q23.3 were extracted from BAM files using samtools. The unmapped sequences were manually aligned to the ERVK11q23.3 DNA sequence using UGENE. The protscale expasy webtool (https://web.expasy.org/protscale/) with a window size of 9 was used to examine the “Hydrophobicity (Kyte & Doolittle)”, “Transmembrane tendency”, and “% buried residues” across the predicted ERVK11q23.3 envelope protein. The ERVK11q23.3 envelope protein sequence containing ORF1 + ORF2 (partial) was used as input; ORF1: MVTPVTWMDNPIEVYVNDSVRVPGPTDDRCPIKPEEEGIMINISTGYRYPICLGRAPGCLIHAVQNWLVEVPTVSPNGRFTYHMVSGMSLRPRVNYLQDFSYQRSLKFRPKGKPCPKEIPKESKNTEVLVWEECVANSAVILQNNEFGTIIDWAPRGQFDHNCSGQTQLCPSAQVSPAVDSDLTESLDKHKHKKLQSLYPWEWGEKGISTPRPKIISPVSGPEHPELWRLIVASHHIRIWSGNQTSETRDRKPFYTIDLNSSLTVPLQSCVKPPYMLVVGNIVIKPDSQTITCENCRLFTCIDSTFNWQQRILLVRAREGVWIPVSMDRPWEASPSIHILTEVLKGILNRSKRFIFTLIAVIMGLIAVTATAAVAGVALHSSVQSVNFVNDWQKNSARLWNSQSSIDQKLANQINHLRQTHLDRRQTHELRTSFPVTV; ORF2(partial): CNTSDFCITPQIYNESEHHWDMVRHHLQGREDNLTLDISKLKEKIFEASKAHLNLVPGTEAIAGVADGLANLNPVTWVKTIGSTTIINLILILVCLFCLLLVCRCTQQLRRDSDHREWAMMTMAVLSKRKGGNVGKSKRDQIVTVSV.

### Cell culture

BeWo cells were cultured at 37°C 5% CO_2_ in Ham’s F-12 (Kaighn’s Modification) media (Caisson Labs, HFL06-500ML) supplemented with 10% FBS (Sigma, 12106C-100ML) and Pen/Strep (Fisher, 15-140-148). Media was changed every other day, and the cells passaged approximately every 4 days or at ∼80% confluency. For fusion induction, BeWo cells were seeded at 20,000 cells/cm^2^, treated with 25uM Forskolin (EMD Millipore, 344282-5MG) one day after passaging, and analyzed 72 h after start of treatment. DMSO treated cells were included alongside forskolin treated cells as negative/vehicle controls.

### siRNA transfection

Custom duplex siRNAs targeting ERVK envelope (siENK) transcripts were designed with IDT’s online siRNA design tool (https://www.idtdna.com/site/order/designtool/index/DSIRNA_CUSTOM), using ERVK11q23.3 sequence as input. In total, three siRNAs targeting ERVK with no predicted cross-reacting transcripts were used: siENK13.7, siENK13.13, and siENK13.34 (Design IDs: CD.Ri.218416.13.7, CD.Ri.218416.13.13, CD.Ri.218416.13.34, respectively); siENK13.13 was predicted to uniquely target ERVK11q23.3 transcripts, while siENK13.7 and siENK13.34 were predicted to target putative ERVK transcripts from 29 and 33 ERVK genomic loci, respectively. For BeWo cells, transfections were performed 24 h after the start of forskolin treatment according to the Lipofectamine RNAiMAX reagent protocol optimized for efficiency, viability and reproducibility. Briefly, for each well of 24-well plate, 1.5μl of RNAiMAX solution (Thermo Fisher, 13778030) and siRNA were diluted in 50μl Opti-MEM media (Thermo Fisher, 31985062), incubated for 5 min at RT, and added to cells containing 500μl culture media (from previous day/containing forskolin). All siRNAs, including the HPRT1 positive control (IDT, 51-01-08-02), nontargeting universal negative control (NC1) (IDT, 51-01-14-03), and ERVK-targeting siRNAs were transfected at a final concentration of 25nM and analyzed 48 h post-transfection (72 h post-forskolin treatment). Fluorescent TYE 563 transfection control was initially used to calculate transfection efficiency at a final concentration of 10nM and analyzed 24 h post-transfection in unstimulated BeWo cells. For HPTs, RNAiMAX reverse transfection protocol was used. For this, siRNA + RNAiMAX complexes were prepared inside the wells, after which the freshly thawed HPT cells and medium were added. All siRNAs, including HPRT1 positive control, NC1, and ERVK-targeting siRNAs were transfected at a final concentration of 25nM and analyzed 72 h post-transfection of HPTs.

### RNA isolation and purification

Frozen placental samples were ground into a powder using a liquid nitrogen-cooled mortar and pestle then directly added to TRIzol reagent (Thermo Fisher #15596026); for cell lines, media was removed and TRIzol reagent was added directly to the tissue culture dish. RNA was isolated from TRIzol reagent, treated with Turbo DNAse (Thermo Fisher #AM1907), and purified using RNA Clean and Concentrator-5 spin columns (Zymo #R1013) according to the manufacturer’s instructions.

### qRT-PCR

qRT-PCR analysis was preformed using qbase+ software, with GAPDH as reference gene and unpaired t-tests (two-sided) used to determine significance. Two sets of gene expression primers, ENK11q23.3_1 and ENK11q23.3_2, were designed to uniquely amplify the ERVK11q23.3 envelope transcript; the first, ENK11q23.3_1, targeted the region of the transcript encoding the surface unit (SU), and ENK11q23.3_2 targeted the region encoding the transmembrane (TM) portion of the envelope protein. An additional primer set, ENK-consensus, was designed to amplify putative ERVK envelope transcript sequences from 17 different genomic loci, including ERVK11q23.3 (chr1:155627693-155627856, chr1:160698826+160698989, chr1:75382268+75382431, chr11:118722041-118722204, chr12:58328486-58328649, chr19:27638644-27638807, chr19:35572497-35572660, chr2:129962985-129963148, chr22:18946662+18946825, chr3:101699824+101699987, chr3:113025296-113025459, chr3:125898575+125898733, chr5:156658733-156658896, chr6:77717964-77718127, chr7:4583453-4583616, chr7:4591957-4592120, chr8:139460933-139461096, chr8:7498902-7499065).

### IF staining

Cell culture media was removed from cells, fixed with ice-cold methanol for 15 min at -20°C, then washed in three changes of PBS. Nonspecific protein binding was blocked by incubating cells in 5% donkey serum for 30 min. Both Anti-ERVK-env mouse monoclonal (HERM-1811-5, Austral Biologicals, San Ramon, CA, USA) and anti-CDH1 rabbit monoclonal (Cell Signaling, 3195S) antibody were diluted 1:250 in PBS + 1% BSA, and incubated overnight at 4°C. Donkey anti-mouse Alexa Fluor 488 (A-21202, Invitrogen) and Donkey anti-rabbit Alexa Fluor 594 (A-21207, Invitrogen) secondary antibodies were diluted 1:1000 in PBS + 1% BSA. The cells were washed in three changes of PBS, before incubating in secondary antibody dilutions for 1 h at RT. Nuclei were counterstained with DAPI and washed with three changes of PBS before imaging.

### Trophoblast cell fusion quantification

A total of five E-cadherin (CDH1) immunostained micrographs were captured from each well using the 20X objective on a Nikon epifluorescence microscope. The CDH1-immunostained micrographs were deidentified and nuclei quantified by a blind reviewer using the “cell counter” tools from the FIJI software package. DAPI staining was used to count the total number of nuclei per micrograph (n_t_). An overlay of CDH1 and DAPI was used to quantify the total number of nuclei within multinucleated cells (n_m_). A multinucleated cell was identified by the presence of two or more nuclei bound by a single membrane of CDH1-positive staining. The fusion percentage was calculated for each micrograph using the following formula.

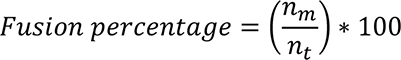

The mean average fusion percentage was calculated for each well, and these values were used to compare fusion levels between siNC1 (n=3) and siENK (n=3) transfected cells using a two-sided unpaired t-test.

## Authors’ contributions

J.L.R, S.L.C. and L.C. designed research; J.L.R performed research; J.L.R., M.M, A.M. analyzed data;

S.L.C. and L.C. supervised progress of research; and J.L.R wrote the manuscript. S.L.C. and L.C. provided edits/comments in the revision and all authors approve of the revised manuscript.

## Acknowledgements

We thank Drs. T. Morgan for providing FFPE placental samples used for IHC; L. Myatt for help obtaining frozen human placental samples used for RNA-seq; A. Valent for providing frozen vials of HPTs; A. Frias for use of tissue culture facilities; A. Adey and A. Fields for the assistance and use of NextSeq500 sequencer; J. Hennebold and M. Murphy for use of their microscope facilities; S. Stadler, A. Adey, J. Sacha for insight and advice.

